# DIANA: Deep Learning Identification and Assessment of Ancient DNA

**DOI:** 10.64898/2026.04.09.717429

**Authors:** Camila Duitama González, Maria Lopopolo, Luca Nishimura, Roland Faure, Sebastian Duchene

## Abstract

The field of ancient metagenomics provides insights into past microbiomes, but with a growing dataset size, methods that rely on reference databases have limited scope. Here, we introduce DIANA, a multi-task neural network that predicts key metadata categories from unitig abundances. Trained on 2,597 run accessions (1.72 Tbp of assembled unitig sequences), DIANA accurately identifies sample host (94.6%), community type (90.0%), and material (88.9%) on held-out test data and demonstrates robust generalisation on an independent validation set. A key innovation is DIANA’s ability to perform semantic generalisation, correctly classifying samples with labels unseen during training — such as novel subspecies — to their appropriate parent categories. By leveraging both known and uncharacterized genomic sequences, DIANA provides a rapid, data-driven system for metadata validation and quality control, accelerating discovery in ancient metagenomics research.

## 1 Introduction

Ancient metagenomics datasets are growing rapidly, with the AncientMetagenomeDir (v25.12.0) [1] now listing 2,980 host-associated and 28 environmental libraries from 216 publications, totalling 6.6 TB of raw data. Despite this wealth of publicly available information, much of it remains underutilised in standard ancient DNA (aDNA) processing pipelines. Traditional reference-based methods—such as read mapping, microbial source tracking, and taxonomic profiling—are computationally prohibitive at this scale and susceptible to reference bias [1–4]. Additionally, these approaches often require time-consuming downloads and local storage of raw data.

Rapid, accurate characterisation of aDNA requires connecting genomic sequences to reliable metadata for each ancient sample. Ideally, researchers could instantly compare new samples with curated datasets (such as AncientMetagenomeDir) to uncover metadata errors, sample mix-ups, contamination, or low-quality data—streamlining downstream analyses.

Directly comparing a new sample to the full 6.6 TB of AncientMetagenomeDir data was nearly impossible until recently. Prior efforts aggregated *k*-mer^1^ counts across collections [5–7], generating massive and redundant *k*-mer matrices. These matrices are not only computationally expensive to produce and store, but also suffer from the “curse of dimensionality” [8] (*p* ≫*n*). Tools like decOM [9] are ultimately limited by the bottleneck of *k*-mer matrix construction [5–7, 10].

Logan has transformed this landscape by making unitig sequences publicly available for every Sequence Read Archive (SRA) run accession [11]. Unitigs offer a compact and exhaustive alternative to raw data, enabling efficient exploration of genomic diversity at scale [12–14]. Constructed via de Bruijn Graph (dBG) assembly, unitigs are the result of collapsing overlapping *k*-mers into single, non-branching paths in the de Bruijn graph (see Figure 1) [15–17]. Tools like MUSET [18] now build matrices tracking unitig presence across thousands of samples. We further advance this by constructing abundance unitig matrices that transform complex genomic data into machine-learning-ready arrays. By integrating these with curated metadata from AncientMetagenomeDir, we can directly link metagenomic content to sample traits.

**Fig. 1.**
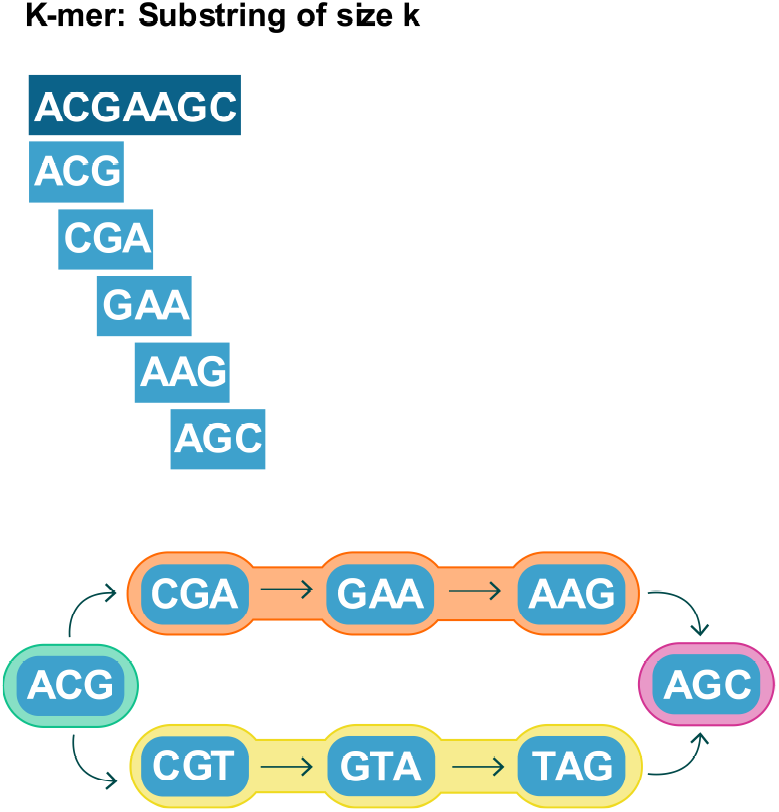
Conceptual illustration of unitig derivation. A DNA sequence (dark blue) is decomposed into its constituent *k*-mers (here, k=3 in lighter blue), which form the nodes of a de Bruijn graph. A unitig is defined as a maximal non-branching path in this graph. The paths highlighted in red, orange, green and yellow are two such unitigs, formed by compacting linear chains of *k*-mers. Critically, nodes that represent branching points, such as “ACG” (green) and “AGC” (pink), are themselves considered maximal non-branching paths of length one and are therefore also unitigs in this set.

On the other hand, recent advances in Deep Learning (DL) have targeted various aDNA tasks, including sample authentication using physicochemical features [19], summary statistics such as GC-content [20], improved variant calling [21], and imputation of genotype probabilities to perform identity-by-descent detection [22]. Some methods directly analyse genomic sequences with CNNs for identification of introgressed fragments or to predict DNA methylation patterns [22, 23].

Despite this progress, no current method can scalably compare a new sample against the full AncientMetagenomeDir dataset. State-of-the-art profilers—KrakenUniq, MetaPhlan, HOPS, aMeta, and other source tracking tools—require thousands of CPU-hours and terabytes of storage just to begin [24–29]. Scalable aDNA characterisation using unitig abundance profiles remains an open challenge.

In this work, we introduce DIANA, an Deep Learning framework for predicting sample metadata from unitig-based metagenomic profiles. For the first time, DIANA enables researchers to comprehensively compare any query metagenomic sample—including ancient samples—against the complete corpus of publicly available data in Logan, enriched with curated metadata from the AncientMetagenomeDir. This approach not only accelerates the assessment of metadata consistency and integrity but also flags quality-control issues (such as contamination or sample mix-ups) and characterises samples with incomplete records, paving the way for more reliable and scalable ancient metagenomic analyses.

## 3 Results

Figure 2 illustrates the DIANA workflow: a multi-task deep learning model trained on 2,597 metagenomic run accessions from AncientMetagenomeDir and Logan, representing each sample as a 107,480-dimensional unitig abundance vector to simultaneously predict Sample Type, Community Type, Sample Host, and Material. We evaluated DIANA on a held-out test set (461 samples) and an independent external validation set (987 samples, including unseen labels). The model achieved high accuracy across all four classification tasks using unitig abundance; that is, the input features of the model were the unitig abundances, evaluated on a training set, a test set, and a validation set of distinct run accessions (samples). See Figure 2.

**Fig. 2.**
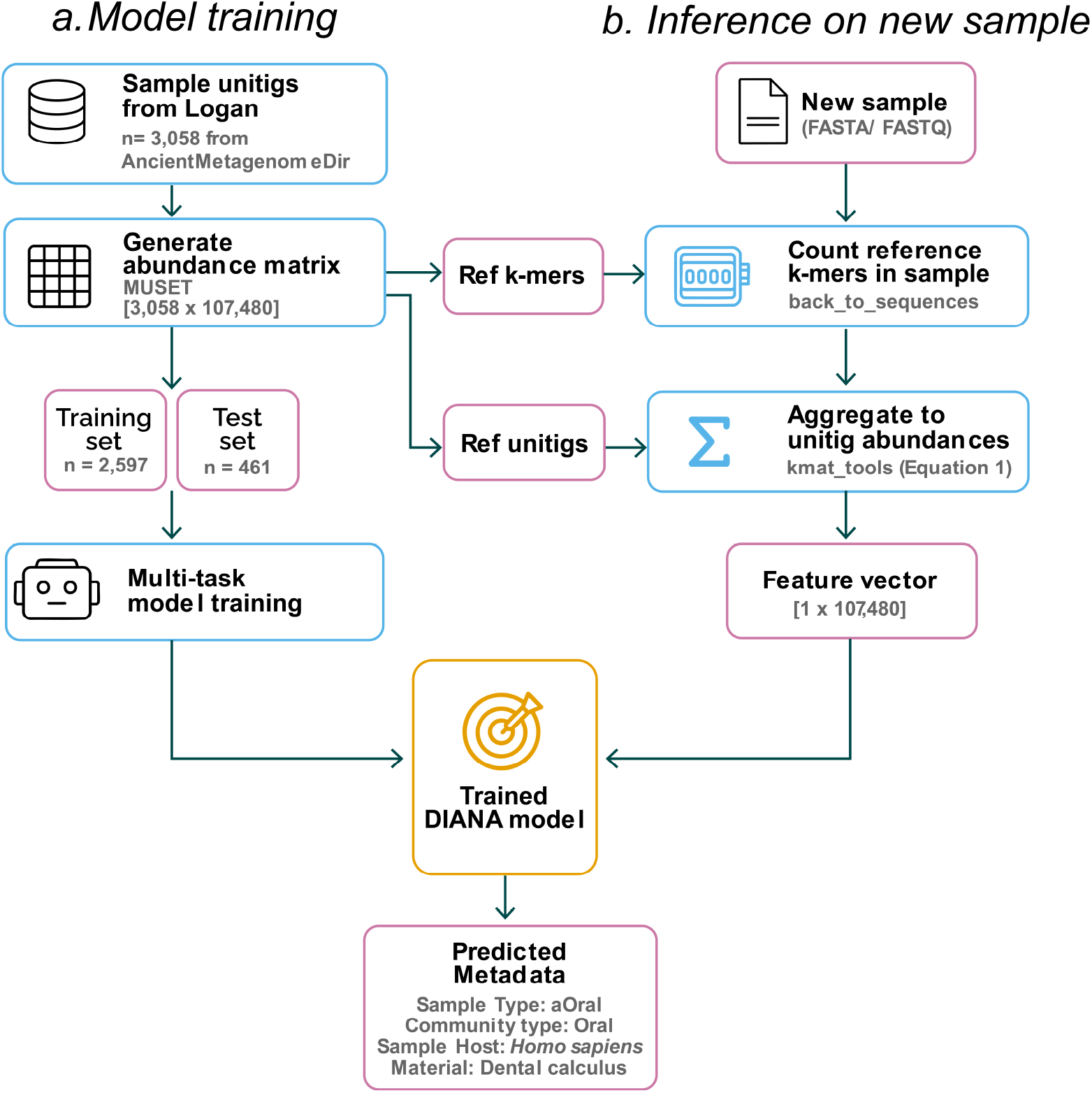
Overview of the DIANA workflow. (a) Model training: Unitig files from samples (run accessions) in aMetagenomeDir and Logan are used to construct a unitig abundance matrix (3,058 samples × 107,480 features), which is partitioned into training (n=2,597) and test (n=461) sets for multi-task neural network training. ((b) Inference pipeline for new samples: Two reference files derived from the training matrix are required: the pre-extracted *k*-mers (`reference_kmers.fasta`; 18.9 million sequences and the unitig sequences (`unitigs.fa`; 107,480 sequences). Raw sequencing reads (FASTQ) from a new sample are queried against `reference_ kmers.fasta` using `back_to_sequences` to count how many reference_*k*-mers are present (minimum abundance of 2 to suppress sequencing errors). `kmat_tools` then uses `unitigs.fa` to map those per-*k*-mer counts to their parent unitigs and computes the fractional abundance following Equation 1, yielding a 107,480-dimensional feature vector. This vector is fed into the trained DIANA model to predict Sample Type, Community Type, Sample Host, and Material.

### 2.1 Robust performance on held-out test data

On the test set, DIANA achieved 99.6% accuracy for Sample Type, 90.0% for Community Type, 94.6% for Sample Host, and 88.9% for Material (Table 1).

**Table 1.**
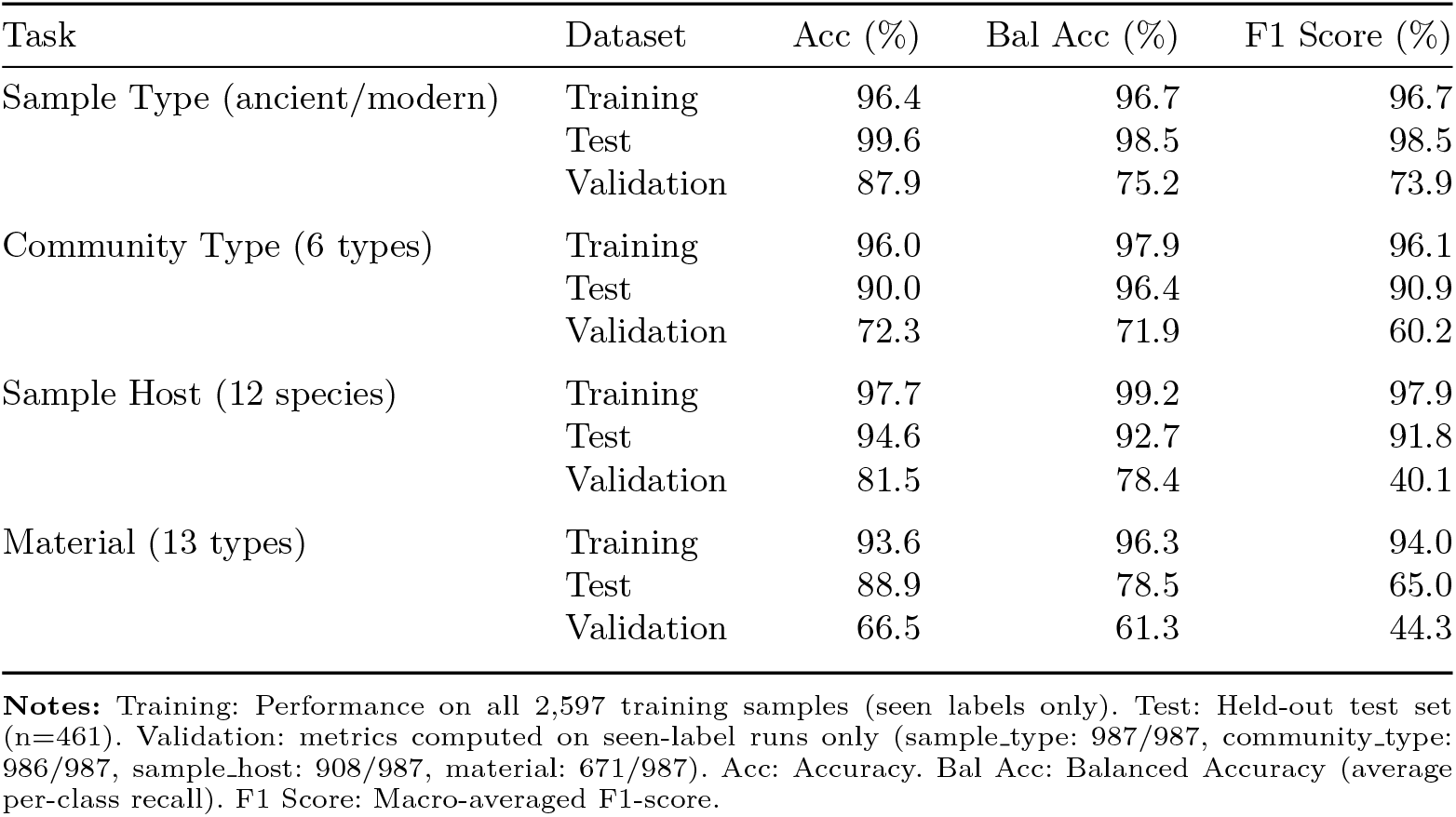
Final model performance across the Training set, the held-out Test set, and the external Validation set. Balanced Accuracy and F1 Score are macro-averaged (unweighted mean across classes), giving equal weight to each class regardless of frequency.

### 2.2 Robust generalisation to an external validation set

On the external validation set, DIANA achieved 87.9% accuracy for distinguishing ancient from modern samples, 72.3% for Community Type, 81.5% for Sample Host, and 66.5% for Material. Detailed per-class results are provided in Supplementary Tables 2 and 4.

Errors in DIANA were concentrated on a few BioProjects and fall into three categories: (i) absent training labels (e.g. birch pitch, intestine), (ii) label granularity where the true label is a subtype of a training class (e.g. lake/marine sediment predicted as sediment, subspecies predicted at genus level), and (iii) genuine confusion between biologically similar classes previously seen (e.g. tooth predicted as dental calculus, oral predicted as skeletal tissue; see Supplementary Table 9).

### 2.3 DIANA learns from biologically informative features

We annotated all 107,480 unitig features using NCBI BLAST. Of these, 77.1% had high-quality BLAST hits (mean sequence identity 98.1%, mean query coverage 98.3%; Supplementary Table 5). The remaining 22.9% (24, 627 unitigs) lacked NCBI hits, but 99.9% (24, 596) matched at least one SRA run in the Logan database with a mean query coverage of 99.7%, mainly from human oral metagenomes and genomic studies (Supplementary Figures 8 and 9). Thus, nearly all DIANA input features correspond to genuine biological sequences.

Additionally, the top-ranked classification features, characterised by taxonomic annotation, are shown in Supplementary Figure 7. These sequences vary systematically across sample types. As expected, many of those sequences correspond to bacterial hits.

Mapping 39 common Illumina adapter sequences against all 107,480 unitigs using Bowtie 2 (v2.4.5) yielded no valid alignments, confirming that the feature set is free of adapter contamination.

### 2.4 Efficient computational performance

DIANA demonstrates fast prediction times. On the validation set (987 samples), runtime shows a moderate positive correlation with input FASTQ file size (*R*^2^ = 0.54; Supplementary Figure 1), averaging approximately 1.8 minutes per GB of input data (Table 2).

**Table 2.** Validation inference computational resources stratified by memory tier (987 samples)

| Memory (GB) | CPUs | N | Runtime (min) | Input size (GB) |
| --- | --- | --- | --- | --- |
| 31 | 6 | 686 | $1.98 \pm 1.52$ | $0.99 \pm 2.20$ |
| 62 | 6 | 130 | $5.10 \pm 2.11$ | $1.40 \pm 0.66$ |
| 125 | 6 | 90 | $11.12 \pm 3.52$ | $3.92 \pm 1.46$ |
| 250 | 6 | 69 | $18.46 \pm 7.36$ | $5.95 \pm 1.89$ |
| 500 | 6 | 12 | $35.89 \pm 17.36$ | $10.78 \pm 3.04$ |

Runtime includes the creation of a vector of unitig feature abundances per sample and neural network inference.

### 2.5 Data Availability: Unitig signature and BLAST annotations

To facilitate reproducibility and enable future research, we publicly release all 107,480 unitig sequences used as features in DIANA, along with comprehensive BLAST annotations against the NCBI nucleotide database (nt) (data deposited on Zenodo under DOI: https://zenodo.org/records/18496961)

### 2.6 DIANA outperforms linear baseline classifiers

No other published tool predicts metadata from unitig abundances, so we compared DIANA to three linear classifiers: Logistic Regression, Linear SVM, and Ridge Classifier. DIANA achieved higher balanced accuracy than all baselines on three of four tasks when evaluated on the training set: Sample Type (0.969 vs. 0.965–0.973), Community Type (0.918 vs. 0.906–0.937), and Sample Host (0.904 vs. 0.836–0.871; Supplementary Figure 10). Linear baselines slightly outperformed DIANA on Material (0.838 vs. 0.812), but their differences were within cross-validation variance (±0.095). Crucially, our model delivers competitive or superior results with a single multi-task model, while baselines require four separate models.

## 3 Methods

### 3.1 Dataset curation and feature matrix generation

We intersected run accessions from AncientMetagenomeDir (v25.09.0) [1], a curated repository of ancient metagenomes, with unitigs from the Logan project [11]. This yielded 3,058 sequencing runs accessions (referred to as “samples” for simplicity), some of which may derive from the same biological specimen, including both ancient and modern metagenomes.

We trained multi-task models to predict four metadata categories. Three categories were derived directly from AncientMetagenomeDir’s standardised metadata schema [30]^2^, with allowed values validated against controlled vocabularies: (i) *sample host*, the Linnean Latin name of the host organism following NCBI taxonomy; (ii) *community type*, the type of microbial community from the host’s body; and (iii) *material*, the sample type from which DNA was extracted, following UBERON (anatomy) or ENVO (environment) ontologies. The fourth category, (iv) *sample type* (ancient vs modern), was manually annotated to distinguish ancient samples from Ancient-MetagenomeDir from modern metagenomic samples we collected from the decOM project [9] or elsewhere in MgNify [31].

Modern metagenomic samples retrieved from public repositories (SRA/ENA) use heterogeneous material terminology that differs from the AncientMetagenomeDir vocabulary used for ancient samples. To ensure consistent classification across the dataset, we mapped modern material labels to terms aligned with the Ancient-MetagenomeDir. Oral-derived samples (supragingival plaque, subgingival plaque, gingiva, tongue, buccal mucosa) were grouped as ‘plaque’, ‘saliva’, or ‘tissue’ based on sample context; broad categories like ‘soft tissue’ and ‘digestive contents’ were mapped to ‘tissue’ and ‘digestive tract contents’; and environment-specific terms were aligned to AncientMetagenomeDir vocabulary (bone, dental calculus, leaf, sediment, shell, soil, tooth). This standardisation resulted in 14 material classes while maintaining biologically meaningful distinctions (e.g., dental calculus, plaque, and saliva as separate categories).

From the previously mentioned 3,058 samples, we generated a unitig abundance matrix using MUSET (v0.6.1) [18]. To focus on informative features, we kept unitigs present in 10–90% of samples, filtering out both rare and ubiquitous unitigs. The matrix used the `--out-frac` parameter, which calculates the fraction of a unitig’s *k*- mers (*k* = 31) present in each sample, providing a semi-quantitative measure of unitig presence. Formally, for a unitig *u* with *N k*-mers and a sample *S*, the fraction *f* (*u, S*) is:

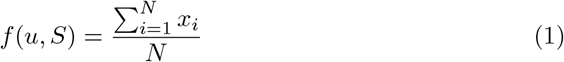

where *x*_*i*_ is a binary indicator of the presence of the *i*-th *k*-mer of *u* in sample *S*. The final feature matrix consisted of **107**,**480 unitig features** across **3**,**058 samples**. Computational resources for matrix generation are detailed in Supplementary Table 8. Principal component analysis revealed that the first 50 components captured only approximately 15% of variance with substantial class overlap in PC1-PC2 space (Supplementary Figures 4 and 5), indicating a non-linear, high-dimensional structure that motivated our use of deep neural networks rather than traditional linear classifiers.

### 3.2 Data partitioning and external validation set

We use three data partitions, each serving a distinct role. The training set (*n* = 2597) and test set (*n* = 461) are both derived from the unitig matrix. The former fits the model and the latter provides a final unbiased performance estimate. The validation set (*n* = 987) was excluded from matrix construction entirely and is processed independently through the dianapredict inference pipeline. Because these samples had no influence on which unitigs were selected as features, the strong performance on this set confirms that the model’s predictive power is not an artefact of feature definition.

The train and test sets were constructed using stratified random sampling with a fixed random seed (42) for reproducibility. Stratification was performed on the “Community type” variable to ensure representative class distributions across splits, with special handling for imbalanced classes. The test set was strictly held out and used only for the final, unbiased evaluation of the trained model. We confirmed that the splits between train and test set were balanced across key metadata variables, including publication year and geographic origin (Supplementary Figures 2 and 3).

To assess real-world performance, we compiled an independent external validation set comprising 987 samples (863 ancient from AncientMetagenomeDir (v25.09.0) and 124 modern metagenomes from MgNify [31]) that were not present in our training or test sets. For these samples, raw FASTQ files were downloaded from the SRA. A full breakdown of sample distributions is available in Supplementary Table 1.

### 3.3 Model architecture and training

In machine learning, a feature is an individual, measurable property of the data used for prediction [8]. For the DIANA model, each metagenomic sample is represented by a feature vector, with each element corresponding to a specific unitig in our reference set of 107,480. The value of each feature is the fractional abundance of its corresponding unitig within that sample (Equation 1). This process transforms the raw sequencing data into a vector, allowing the model to classify samples based on their distinct unitig abundance profiles.

DIANA employs a multi-task feedforward neural network that simultaneously predicts all four metadata categories (Sample Type, Community Type, Sample Host, and Material) from the shared unitig feature vector. The architecture consists of an input layer accepting 107,480 features, followed by two fully connected hidden layers (269 and 371 neurons) with ReLU activation and dropout regularisation (rate = 0.1696) to prevent overfitting. The network then branches into four task-specific output heads, each terminating in a softmax layer to produce class probabilities. The final optimised hyperparameters are detailed in Supplementary Table 3.

Hyperparameter tuning was performed on the training set using a nested cross-validation scheme (5-fold outer loop, 3-fold inner loop) with 50 Optuna [32] trials per fold. The final model was trained on 90% of the full training set (n=2,337), with the remaining 10% (n=260) used as an internal validation set for early stopping.

### 3.4 Model evaluation and feature analysis

The final model was evaluated on the held-out test set. We computed standard classification metrics for each task, including accuracy, balanced accuracy and F1-score. Reciever Operating Characteristic (ROC) and Precision-Recall (PR) curves were generated to assess discriminative performance.

To contextualise DIANA’s performance, we compared it against three linear baseline classifiers on the training set — Logistic Regression, Linear Support Vector Machine, and Ridge Classifier — each trained independently per task with class-balanced weighting, using the same 5-fold cross-validation scheme and balanced accuracy as the primary metric.

To identify the most influential features, we applied gradient-based feature attribution. The top 100 most discriminant unitigs for each task were selected and taxonomically annotated using BLAST+ v2.16.0 against the NCBI nucleotide database (E-value ≤ 1 × 10^*−*5^). Taxonomy was assigned using a hierarchical approach: genus-level classification when available, otherwise family-level, and finally phylum-level. DIANA’s unitigs that did not have a blast hit were posteriously aligned using LexicMap[33] to all the unitigs of Logan flagged as “bacterial genomic”[11].

Finally, to verify the biological integrity of the feature set, we mapped 39 curated Illumina adapter sequences against all 107,480 unitigs using Bowtie 2 (v2.4.5) [34]; full details are provided in Supplementary Section 3.1.

### 3.5 The dianapredict pipeline for new samples

For the external validation set, and for general use on new query samples, we developed the `dianapredict` pipeline to reconstruct the unitig feature vector directly from raw sequencing reads (FASTA/FASTQ files). The 107,480 training unitigs define the reference vocabulary: their constituent *k*-mers (*k* = 31) are pre-extracted from the MUSET matrix and stored as `reference_kmers.fasta` (18,953,119 sequences of 31 bp). This three-step process involves: (1) counting how many of these reference *k*- mers are present in the new sample using `back_to_sequences` [35], with a minimum abundance of 2 to filter sequencing errors; (2) aggregating per-*k*-mer counts to unitig-level fractions and (3) feeding the resulting 107,480-dimensional feature vector into the trained DIANA model for classification.

## 4 Discussion

We introduced DIANA, the first Deep Learning framework to classify ancient metagenomic samples using unitig abundance as features. Our results demonstrate that unitig profiles provide strong, discriminative signals for predicting key metadata categories with high accuracy, offering a valuable tool for the paleometagenomics community.

While DIANA does not aim at replacing standard taxonomic profilers and authentication pipelines, it offers a first approach at comparing a new aDNA sample against all the samples in the AncientMetagenomeDir, in a way that no other taxonomic classifier or pipeline could currently achieve. Constructing taxonomic profiles for the full corpus using standard profilers such as aMeta [27], KrakenUniq [24], or MetaPhlAn[25] would require an estimated thousands to millions of CPU-hours, depending on the tool, and must be completed before any cross-sample comparison can begin. Moreover, all of these approaches require first downloading and locally storing the complete 6.6 TB of raw FASTQ files from all the run accessions (samples) used by our model. Instead, DIANA compares any new aDNA sample against all publicly available ancient metagenomic samples in **under 2 minutes** for the majority of samples, on 6 CPUs and as little as 31 GB of RAM, requiring only the sample FASTQ file and a 750 MB reference *k*-mer file.

### 4.1 DIANA as a tool for metadata validation and anomaly detection

DIANA’s primary role is to serve as a **metadata validation and contextualization tool**. In large-scale aDNA studies, sample mix-ups and mislabeling are persistent challenges that can compromise downstream results. By comparing a sample’s genomic profile to thousands of known references, DIANA provides an independent, data-driven assessment of its biological content, flagging inconsistencies that may indicate such errors. To be clear, DIANA does not evaluate DNA damage patterns or fragment length distributions, which remain among the main methods for authenticating ancient molecules. Instead, it complements these methods by ensuring the sample’s reported metadata (e.g., host, material, community type) is consistent with its sequencing data.

### 4.2 Learning biologically and semantically meaningful patterns

Feature importance analysis confirmed DIANA’s predictions are driven by taxonomically informative unitigs, such as oral microbes for “dental calculus” samples (Supplementary Figure 7). The fact that nearly half of these highly informative unitigs lack matches in public databases suggests that DIANA leverages novel genomic signals inaccessible to standard reference-based methods (Supplementary Table 5). Notably, DIANA makes remarkably insightful predictions even for categories it was never trained on. For example, it correctly maps novel primate subspecies such as *Gorilla beringei* to the more general *Gorilla sp*. class learned from the training data. Similarly, it classifies unseen materials such as “lake sediment” and “marine sediment” under the broader “sediment” category, demonstrating a capacity for both phylogenetically and semantically informed “zero-shot” generalisation (Supplementary Table 6). This makes DIANA a powerful tool for generating hypotheses about uncharacterized samples.

Moreover, we mapped 39 common Illumina adapter sequences against all the unitigs that Diana uses as features, using Bowtie 2 (v2.4.5). This analysis yielded no valid alignments (see Supplementary Section 1), ruling out spurious correlations due to laboratory or protocol-specific adapter usage. These results confirm that the class-discriminative signal learned by Diana is biological in origin.

### 4.3 Limitations and future directions

DIANA has two main limitations. First, as a supervised classifier, it achieves lower accuracy on rare classes with limited training data, such as “soft tissue” (Supplementary Table 2). This challenge is inherent but is expected to diminish as curated datasets expand and underrepresented classes gain more examples. Additionally, increasing the availability of well-annotated modern metagenomes—akin to the AncientMetagenomeDir for ancient samples—would further enhance the comparison and contextualization of ancient metagenomic data.

In the external validation set, errors are concentrated in a small number of BioProjects, reflecting three distinct causes: absent training labels, label granularity, and genuine confusion between similar classes (Supplementary Table 9).

Certain decisions made during the initial unitig matrix construction, such as *k*mer size, minimum abundance, minimizer size, and others (see Supplementary Table 7), should be optimised to assess their impact on the model’s performance. However, this was not done for this manuscript, as constructing the unitig matrix remains computationally challenging given the size of the input dataset.

Finally, the model’s accuracy could be further improved by enriching the feature set. Incorporating classic aDNA signals, such as damage patterns and fragment-length statistics, would provide orthogonal information to unitig profiles and likely enhance the model’s performance, especially for distinguishing ancient from modern samples. Furthermore, as the AncientMetagenomeDir metadata is further curated and includes more samples, DIANA could be updated and improve its predictive capacity.

## 5 Conclusion

DIANA is a powerful tool that should be included in ancient metagenomic authentication pipelines. It offers a rapid and scalable way to contextualise new samples within the landscape of published aDNA data. By flagging inconsistencies between metadata and genomic content, DIANA helps researchers identify potential issues early, validate sample integrity, and generate robust hypotheses about unknown samples. As the scale of paleogenomics continues to expand, tools like DIANA will be essential for maintaining data quality and accelerating discovery.

## Supplementary information

Supplementary tables and figures are in the accompanying PDF file.

## Acknowledgements

The authors thank the anonymous reviewers for their valuable suggestions. We thank the SPAAM community and the AncientMetagenome Core team, which maintains the repository through regular releases. We also thank Manuela Aguirre Botero, Gaston Rijo de León and Rayan Chikhi for their input on this manuscript. Finally, we thank Camila Bolívar Manzano for her help designing the illustrations for DIANA.

## Declarations

### Ethics approval and consent to participate

Not applicable. This study was conducted using publicly available data from the Sequence Read Archive. No new samples were collected, and no experiments involving human participants or animals were performed.

### Consent for publication

Not applicable. This study does not contain any individual person’s data in any form.

### Data availability

All raw sequencing data used in this study are publicly available from the Sequence Read Archive (SRA). A complete list of SRA accession numbers for the training, test, and validation datasets is available in the project’s GitHub repository: https://github.com/CamilaDuitama/DIANA/tree/paper/paper/accessions. The 107,480 unitig sequences used as features, along with their comprehensive BLAST annotations, have been deposited on Zenodo under DOI: https://zenodo.org/records/18496961.

### Materials availability

Not applicable. This study did not generate new physical materials.

### Code availability

The source code for DIANA is publicly available on GitHub at https://github.com/CamilaDuitama/DIANA. The pre-trained model weights and PCA reference file are hosted on Hugging Face Hub at https://huggingface.co/cduitamag/DIANA and are automatically downloaded by the installer.

### Author contributions

**C.D.G**.: Conceptualisation, Data curation, Formal analysis, Investigation, Methodology, Software, Validation, Visualisation, Writing - original draft. **M.L:** Validation, Writing - Review & Editing. **R.F:** Formal analysis, Validation, Writing. **S.D**.: Conceptualisation, Supervision, Funding acquisition, Writing – Review and Editing. **L.N:** Funding acquisition, Validation, Writing - Review & Editing.

## Abbreviations

**Acronyms**

aDNA: ancient DNA
CNN: Convolutional Neural Network
dBG: de Bruijn Graph
DL: Deep Learning
PR: Precision-Recall
ROC: Reciever Operating Characteristic
SRA: Sequence Read Archive
SVM: Support Vector Machine

## Footnotes

1 substring of a genomic sequence of length *k*

2 https://github.com/SPAAM-community/AncientMetagenomeDir/tree/master/assets/enums

