## Supplementary file for "DIANA: Deep Learning Identification and Assessment of Ancient DNA"

Supplementary Information for:
### DIANA: Deep Learning Identification and Assessment of Ancient DNA

Camila Duitama González et al.

### 1 Supplementary Tables

Supplementary Table 1: Sample distribution across class labels for each prediction task and dataset split

| Task | Class | Training | Test | Validation | Total |
|---|---|---|---|---|---|
| Sample Type | ancient | 2419 | 425 | 863 | 3707 |
| | modern | 178 | 36 | 124 | 338 |
| Community Type | Not applicable - env sample | 615 | 110 | 265 | 990 |
| | gut | 86 | 15 | 37 | 138 |
| | oral | 1460 | 260 | 562 | 2282 |
| | plant tissue | 81 | 14 | 7 | 102 |
| | skeletal tissue | 267 | 47 | 82 | 396 |
| | soft tissue | 88 | 15 | 33 | 136 |
| Sample Host | *Ambrosia artemisiifolia* | 57 | 9 | 3 | 69 |
| | *Arabidopsis thaliana* | 24 | 5 | 4 | 33 |
| | *Canis lupus* | 11 | 8 | 2 | 21 |
| | *Gorilla sp.* | 110 | 25 | 0 | 135 |
| | *Homo sapiens* | 1418 | 230 | 586 | 2234 |
| | *Homo sapiens neanderthalensis* | 25 | 10 | 9 | 44 |
| | *Mammuthus primigenius* | 12 | 4 | 0 | 16 |
| | Not applicable - env sample | 615 | 110 | 265 | 990 |
| | Other mammal | 9 | 3 | 0 | 12 |
| | *Pan troglodytes* | 40 | 8 | 0 | 48 |
| | *Rangifer tarandus* | 19 | 5 | 0 | 24 |
| | *Ursus arctos* | 257 | 44 | 39 | 340 |
| | *Alouatta palliata* | 0 | 0 | 8 | 8 |
| | *Gorilla beringei* | 0 | 0 | 4 | 4 |
| | *Gorilla beringei beringei* | 0 | 0 | 16 | 16 |
| | *Gorilla beringei graueri* | 0 | 0 | 12 | 12 |
| | *Gorilla gorilla gorilla* | 0 | 0 | 5 | 5 |
| | *Homo sapiens* | 0 | 0 | 1 | 1 |
| | *Lithobates pipiens* | 0 | 0 | 11 | 11 |
| | *Pan troglodytes ellioti* | 0 | 0 | 4 | 4 |
| | *Pan troglodytes schweinfurthii* | 0 | 0 | 7 | 7 |
| | *Papio anubis* | 0 | 0 | 4 | 4 |
| | *Papio hamadryas* | 0 | 0 | 5 | 5 |
| | *Papio sp.* | 0 | 0 | 1 | 1 |
| | *Unknown* | 0 | 0 | 1 | 1 |
| | Oral | 0 | 0 | 1 | 1 |
| | Skin | 69 | 11 | 20 | 100 |

*Continued on next page*

*Table 1 continued from previous page*

| Task | Class | Training | Test | Validation | Total |
|---|---|---|---|---|---|
| | anterior nares | 0 | 0 | 2 | 2 |
| | birch pitch | 0 | 0 | 79 | 79 |
| | bone | 195 | 41 | 50 | 286 |
| | brain | 0 | 0 | 1 | 1 |
| | buccal mucosa | 0 | 0 | 2 | 2 |
| | dental calculus | 1130 | 205 | 356 | 1691 |
| | dentine | 0 | 0 | 2 | 2 |
| | digestive_contents | 92 | 17 | 1 | 110 |
| | faeces | 0 | 0 | 17 | 17 |
| | gingiva | 1 | 1 | 0 | 2 |
| | intestine | 0 | 0 | 11 | 11 |
| | lake sediment | 0 | 0 | 91 | 91 |
| | leaf | 81 | 14 | 7 | 102 |
| | marine sediment | 0 | 0 | 66 | 66 |
| | midden | 11 | 1 | 1 | 13 |
| | palaeofaeces | 0 | 0 | 5 | 5 |
| | permafrost | 58 | 13 | 14 | 85 |
| | plaque | 0 | 0 | 19 | 19 |
| | saliva | 9 | 3 | 5 | 17 |
| | sediment | 450 | 78 | 40 | 568 |
| | shallow marine sediment | 0 | 0 | 3 | 3 |
| | shell | 19 | 5 | 0 | 24 |
| | soft_tissue | 13 | 2 | 0 | 15 |
| | soil | 77 | 13 | 53 | 143 |
| | subgingival plaque | 3 | 1 | 0 | 4 |
| | supragingival plaque | 6 | 0 | 0 | 6 |
| | throat swab | 12 | 5 | 0 | 17 |
| | tongue | 1 | 1 | 4 | 6 |
| | tooth | 370 | 50 | 126 | 546 |

**Columns**: *Task*—one of the four prediction tasks (Sample Type, Community Type, Sample Host, Material). *Class*—label category within that task; species names in *Sample Host* are italicised. *Training / Test / Validation*—number of samples carrying that label in each dataset split. *Total*—sum across all three splits. Validation set: 987 samples with successful predictions. Classes with 0 training samples are unseen by the model. Classes with 0 validation samples were present in training but did not appear in the external validation set.

Supplementary Table 2: DIANA predictions for validation samples whose true label was absent from training (unseen classes)

| Task | True Label (Unseen) | Predicted Label | Count |
|---|---|---|---|
| **Sample Host** | | | |
| | *alouatta palliata* | *Gorilla sp.* | 8 |
| | *gorilla beringei* | *Gorilla sp.* | 4 |
| | *gorilla beringei beringei* | *Gorilla sp.* | 16 |
| | *gorilla beringei graueri* | *Gorilla sp.* | 8 |
| | *gorilla beringei graueri* | *Homo sapiens neanderthalensis* | 4 |
| | *gorilla gorilla gorilla* | *Gorilla sp.* | 5 |
| | *homo sapiens* | *Homo sapiens* | 1 |
| | *lithobates pipiens* | *Homo sapiens neanderthalensis* | 11 |
| | *pan troglodytes ellioti* | *Pan troglodytes* | 4 |
| | *pan troglodytes schweinfurthii* | *Pan troglodytes* | 7 |
| | *papio anubis* | *Homo sapiens* | 3 |
| | *papio anubis* | *Pan troglodytes* | 1 |
| | *papio hamadryas* | *Homo sapiens* | 4 |
| | *papio hamadryas* | *Gorilla sp.* | 1 |
| | *papio sp.* | *Homo sapiens neanderthalensis* | 1 |
| | *unknown* | *Homo sapiens neanderthalensis* | 1 |
| **Material** | | | |
| | Oral | tissue | 1 |
| | Skin | skin | 18 |
| | Skin | tooth | 2 |
| | anterior nares | bone | 2 |
| | birch pitch | sediment | 39 |
| | birch pitch | skin | 30 |
| | birch pitch | saliva | 7 |
| | birch pitch | bone | 3 |
| | brain | tissue | 1 |
| | buccal mucosa | skin | 2 |
| | dentine | bone | 1 |
| | dentine | dental calculus | 1 |
| | digestive_contents | digestive tract contents | 1 |
| | faeces | digestive tract contents | 16 |
| | faeces | dental calculus | 1 |
| | intestine | skin | 7 |
| | intestine | bone | 3 |
| | intestine | tooth | 1 |
| | lake sediment | sediment | 81 |
| | lake sediment | permafrost | 9 |
| | lake sediment | tissue | 1 |
| | marine sediment | sediment | 64 |
| | marine sediment | shell | 2 |

*Continued on next page*

*Table 2 continued from previous page*

| Task | True Label (Unseen) | Predicted Label | Count |
|---|---|---|---|
| | palaeofaeces | digestive tract contents | 3 |
| | palaeofaeces | dental calculus | 1 |
| | palaeofaeces | sediment | 1 |
| | shallow marine sediment | sediment | 2 |
| | shallow marine sediment | skin | 1 |
| | tongue | skin | 2 |
| | tongue | tissue | 2 |

**Columns**: *Task*—prediction task in which the unseen label occurs. *True Label (Unseen)*—ground-truth class not present in the training set. *Predicted Label*—class assigned by DIANA (always a seen training class). *Count*—number of validation samples with that true $\rightarrow$ predicted pair. Sample Host: Novel species or subspecies are mapped by DIANA to the nearest genus- or species-level class seen in training. Material: Novel material types are mapped to the semantically closest training class. Sample Host: 79 unseen predictions. Material: 305 unseen predictions.

Supplementary Table 3: Per-class performance on validation set (seen labels only)

| Task | Class Label | n | Accuracy | Precision | Recall | F1-Score |
|---|---|---|---|---|---|---|
| **Sample Type** | | | | | | |
| | ancient | 863 | 92.2% | 93.9% | 92.2% | 93.0% |
| | modern | 124 | 58.1% | 51.8% | 58.1% | 54.8% |
| **Community Type** | | | | | | |
| | oral | 562 | 64.2% | 98.6% | 64.2% | 77.8% |
| | Not applicable - env sample | 265 | 90.9% | 79.8% | 90.9% | 85.0% |
| | skeletal tissue | 82 | 81.7% | 38.1% | 81.7% | 51.9% |
| | gut | 37 | 54.1% | 83.3% | 54.1% | 65.6% |
| | soft tissue | 33 | 54.5% | 17.5% | 54.5% | 26.5% |
| | plant tissue | 7 | 85.7% | 40.0% | 85.7% | 54.5% |
| **Sample Host** | | | | | | |
| | *Homo sapiens* | 586 | 75.8% | 97.6% | 75.8% | 85.3% |
| | Not applicable - env sample | 265 | 92.5% | 77.8% | 92.5% | 84.5% |
| | *Ursus arctos* | 39 | 100.0% | 97.5% | 100.0% | 98.7% |
| | *Homo sapiens neanderthalensis* | 9 | 55.6% | 27.8% | 55.6% | 37.0% |
| | *Arabidopsis thaliana* | 4 | 75.0% | 33.3% | 75.0% | 46.2% |
| | *Ambrosia artemisiifolia* | 3 | 100.0% | 60.0% | 100.0% | 75.0% |
| | *Canis lupus* | 2 | 50.0% | 8.3% | 50.0% | 14.3% |
| **Material** | | | | | | |
| | dental calculus | 356 | 84.6% | 86.5% | 84.6% | 85.5% |
| | tooth | 126 | 23.8% | 40.5% | 23.8% | 30.0% |
| | soil | 53 | 37.7% | 76.9% | 37.7% | 50.6% |
| | bone | 50 | 52.0% | 40.0% | 52.0% | 45.2% |
| | sediment | 40 | 100.0% | 53.3% | 100.0% | 69.6% |
| | plaque | 19 | 78.9% | 65.2% | 78.9% | 71.4% |
| | permafrost | 14 | 50.0% | 87.5% | 50.0% | 63.6% |
| | leaf | 7 | 85.7% | 46.2% | 85.7% | 60.0% |
| | saliva | 5 | 0.0% | 0.0% | 0.0% | 0.0% |
| | midden | 1 | 100.0% | 100.0% | 100.0% | 100.0% |

Metrics computed only for seen labels (present in training set). Accuracy: proportion of samples in each class correctly classified. Precision: proportion of predictions for a class that were correct. Recall: proportion of true instances of a class that were correctly predicted. F1-Score: harmonic mean of precision and recall.

Supplementary Table 4: Optimized hyperparameters for the DIANA multi-task neural network

| Category | Parameter | Value |
|---|---|---|
| Architecture | Input features | 107,480 |
| | Hidden layers | [269, 371] |
| | Dropout rate | 0.1696 |
| | Activation | relu |
| Training | Learning rate | 0.002447 |
| | Weight decay | 2.92e-04 |
| | Batch size | 96 |
| | Max epochs | 200 |
| Task Weights | Sample Type | 1.362 |
| | Community Type | 1.237 |
| | Sample Host | 1.585 |
| | Material | 1.357 |

**Columns**: *Category*—group of related parameters: *Architecture* (network structure), *Training* (optimisation settings), *Task Weights* (per-task loss weight in the joint objective). *Parameter*—hyperparameter name. *Value*—value used in the final model. Hyperparameters were selected by Optuna (50 trials, 5-fold cross-validation); the final model was then trained from scratch on the full training set using these best-found values. Input features: 107,480 unitigs. Max epochs: 200 with early stopping (patience = 20) on validation loss.

Supplementary Table 5: Misclassification patterns on the validation set for seen classes

| Task | True Label | Predicted Label | Count |
|---|---|---|---|
| Sample Type | ancient | modern | 67 |
| | modern | ancient | 52 |
| Community Type | oral | skeletal tissue | 90 |
| | oral | soft tissue | 55 |
| | oral | Not applicable - env sample | 52 |
| | soft tissue | skeletal tissue | 15 |
| | Not applicable - env sample | soft tissue | 13 |
| | gut | soft tissue | 9 |
| | Not applicable - env sample | plant tissue | 9 |
| | skeletal tissue | soft tissue | 8 |
| | skeletal tissue | Not applicable - env sample | 4 |
| | oral | gut | 4 |
| | gut | Not applicable - env sample | 4 |
| | skeletal tissue | oral | 3 |
| | gut | oral | 2 |
| | gut | skeletal tissue | 2 |
| | Not applicable - env sample | skeletal tissue | 2 |
| | plant tissue | Not applicable - env sample | 1 |
| Sample Host | *Homo sapiens* | Not applicable - env sample | 65 |
| | *Homo sapiens* | *Gorilla sp.* | 29 |
| | *Homo sapiens* | *Pan troglodytes* | 19 |
| | *Homo sapiens* | *Homo sapiens neanderthalensis* | 13 |
| | *Homo sapiens* | *Canis lupus* | 11 |
| | Not applicable - env sample | *Homo sapiens* | 10 |
| | Not applicable - env sample | *Arabidopsis thaliana* | 6 |
| | *Homo sapiens* | *Mammuthus primigenius* | 3 |
| | *Homo sapiens neanderthalensis* | Not applicable - env sample | 3 |
| | Not applicable - env sample | *Ambrosia artemisiifolia* | 2 |
| | Not applicable - env sample | *Gorilla sp.* | 2 |

*Continued on next page*

*Table 5 continued from previous page*

| Task | True Label | Predicted Label | Count |
|---|---|---|---|
| | *Canis lupus* | Not applicable - env sample | 1 |
| | *Homo sapiens* | Other mammal | 1 |
| | *Arabidopsis thaliana* | Not applicable - env sample | 1 |
| | *Homo sapiens neanderthalensis* | *Homo sapiens* | 1 |
| | *Homo sapiens* | *Ursus arctos* | 1 |
| Material | tooth | dental calculus | 44 |
| | tooth | bone | 30 |
| | dental calculus | tooth | 26 |
| | bone | tooth | 16 |
| | soil | sediment | 16 |
| | soil | skin | 10 |
| | dental calculus | bone | 9 |
| | dental calculus | skin | 7 |
| | tooth | skin | 7 |
| | dental calculus | soil | 6 |
| | tooth | sediment | 5 |
| | soil | leaf | 5 |
| | permafrost | sediment | 5 |
| | saliva | plaque | 4 |
| | bone | sediment | 4 |
| | dental calculus | sediment | 4 |
| | tooth | plaque | 4 |
| | tooth | digestive tract contents | 3 |
| | tooth | shell | 3 |
| | bone | skin | 2 |
| | dental calculus | saliva | 2 |
| | plaque | dental calculus | 2 |
| | permafrost | leaf | 2 |
| | dental calculus | digestive tract contents | 1 |
| | leaf | sediment | 1 |
| | bone | dental calculus | 1 |
| | bone | shell | 1 |
| | saliva | tooth | 1 |
| | soil | permafrost | 1 |
| | plaque | saliva | 1 |
| | plaque | skin | 1 |
| | soil | tooth | 1 |

**Columns**: *Task*—prediction task in which the error occurred (Sample Type, Community Type, Sample Host, Material). *True Label*—ground-truth class, always present in the training set. *Predicted Label*—class assigned by DIANA. *Count*—number of validation samples with that true → predicted pair. Only misclassifications of seen classes (present in training) are included; rows are sorted by frequency within each task (most common errors first). Species names in *Sample Host* are italicised.

Supplementary Table 6: BLAST annotation of DIANA's 107,480 unitig features against the NCBI nt database.

| **Metric** | **Value** |
|---|---|
| *Overall Statistics* | |
| Total features analyzed | 107,480 |
| Features with BLAST hits | 82,853 |
| Overall hit rate | 77.09% |
| *Identity Distribution (best hit per feature)* | |
| ≥95% identity | 73,166 |
| 90–95% identity | 6,358 |
| 80–90% identity | 2,784 |
| <80% identity | 545 |
| *Top 10 most frequent species (best informative hit per feature)* | |
| *Actinomyces sp.* | 19,754 |
| *Olsenella sp.* | 13,374 |
| *Desulfobulbus oralis* | 12,286 |
| *Eubacterium minutum* | 9,284 |
| *Anaerolineaceae bacterium* | 8,419 |
| *Arachnia propionica* | 3,270 |
| *Fretibacterium fastidiosum* | 1,935 |
| *Homo sapiens* | 1,047 |
| *Actinomyces oris* | 981 |
| *Streptococcus sanguinis* | 771 |

Unitig features form the input to DIANA's multi-task classifier; this table summarises their biological annotation via BLAST (E-value $\leq 10^{-5}$) against the NCBI nt database. *Overall Statistics*: number and fraction of features with at least one hit. *Identity Distribution*: per cent nucleotide identity of the single best hit (highest bitscore) per matched feature. *Top 10 most frequent species*: species most often assigned as the best biologically informative hit; non-specific terms (*metagenome*, *uncultured*, etc.) are excluded before ranking.

Supplementary Table 7: Validation set performance: Seen vs unseen labels with top 10 most frequent misclassification patterns

| Task | Category | True Label | Predicted Label | Count | % Task | % Total |
|---|---|---|---|---|---|---|
| Sample Type | Seen - Correct | — | — | 868 | 87.9% | 87.9% |
| | Seen - Wrong | ancient | modern | 67 | 6.8% | 6.8% |
| | | modern | ancient | 52 | 5.3% | 5.3% |
| Community Type | Seen - Correct | — | — | 713 | 72.2% | 72.2% |
| | Seen - Wrong | oral | skeletal tissue | 90 | 9.1% | 9.1% |
| | | oral | soft tissue | 55 | 5.6% | 5.6% |
| | | oral | Not applicable - env sample | 52 | 5.3% | 5.3% |
| | | soft tissue | skeletal tissue | 15 | 1.5% | 1.5% |
| | | Not applicable - env sample | soft tissue | 13 | 1.3% | 1.3% |
| | | gut | soft tissue | 9 | 0.9% | 0.9% |
| | | Not applicable - env sample | plant tissue | 9 | 0.9% | 0.9% |
| | | skeletal tissue | soft tissue | 8 | 0.8% | 0.8% |
| | | skeletal tissue | Not applicable - env sample | 4 | 0.4% | 0.4% |
| | | oral | gut | 4 | 0.4% | 0.4% |
| | Unseen | Unknown | soft tissue | 1 | 0.1% | 0.1% |
| Sample Host | Seen - Correct | — | — | 740 | 75.0% | 75.0% |
| | Seen - Wrong | *Homo sapiens* | Not applicable - env sample | 65 | 6.6% | 6.6% |
| | | *Homo sapiens* | *Gorilla sp.* | 29 | 2.9% | 2.9% |
| | | *Homo sapiens* | *Pan troglodytes* | 19 | 1.9% | 1.9% |
| | | *Homo sapiens* | *H. s. neanderthalensis* | 13 | 1.3% | 1.3% |
| | | *Homo sapiens* | *Canis lupus* | 11 | 1.1% | 1.1% |
| | | Not applicable - env sample | *Homo sapiens* | 10 | 1.0% | 1.0% |
| | | Not applicable - env sample | *Arabidopsis thaliana* | 6 | 0.6% | 0.6% |
| | | *Homo sapiens* | *Mammuthus primigenius* | 3 | 0.3% | 0.3% |
| | | *H. s. neanderthalensis* | Not applicable - env sample | 3 | 0.3% | 0.3% |
| | | Not applicable - env sample | *Ambrosia artemisiifolia* | 2 | 0.2% | 0.2% |
| | Unseen | *gorilla beringei beringei* | *Gorilla sp.* | 16 | 1.6% | 1.6% |
| | | *lithobates pipiens* | *H. s. neanderthalensis* | 11 | 1.1% | 1.1% |
| | | *gorilla beringei graueri* | *Gorilla sp.* | 8 | 0.8% | 0.8% |
| | | *alouatta palliata* | *Gorilla sp.* | 8 | 0.8% | 0.8% |
| | | *pan troglodytes schweinfurthii* | *Pan troglodytes* | 7 | 0.7% | 0.7% |
| | | *gorilla gorilla gorilla* | *Gorilla sp.* | 5 | 0.5% | 0.5% |
| | | *gorilla beringei graueri* | *H. s. neanderthalensis* | 4 | 0.4% | 0.4% |
| | | *gorilla beringei* | *Gorilla sp.* | 4 | 0.4% | 0.4% |
| | | *papio hamadryas* | *Homo sapiens* | 4 | 0.4% | 0.4% |

Continued on next page

Supplementary Table 7 – continued from previous page

| Task | Category | True Label | Predicted Label | Count | % Task | % Total |
|---|---|---|---|---|---|---|
| | | *pan troglodytes ellioti* | *Pan troglodytes* | 4 | 0.4% | 0.4% |
| Material | Seen - Correct | — | — | 446 | 45.2% | 45.2% |
| | Seen - Wrong | tooth | dental calculus | 44 | 4.5% | 4.5% |
| | | tooth | bone | 30 | 3.0% | 3.0% |
| | | dental calculus | tooth | 26 | 2.6% | 2.6% |
| | | bone | tooth | 16 | 1.6% | 1.6% |
| | | soil | sediment | 16 | 1.6% | 1.6% |
| | | soil | skin | 10 | 1.0% | 1.0% |
| | | dental calculus | bone | 9 | 0.9% | 0.9% |
| | | dental calculus | skin | 7 | 0.7% | 0.7% |
| | | tooth | skin | 7 | 0.7% | 0.7% |
| | | dental calculus | soil | 6 | 0.6% | 0.6% |
| | Unseen | lake sediment | sediment | 81 | 8.2% | 8.2% |
| | | marine sediment | sediment | 64 | 6.5% | 6.5% |
| | | birch pitch | sediment | 39 | 4.0% | 4.0% |
| | | birch pitch | skin | 30 | 3.0% | 3.0% |
| | | Skin | skin | 18 | 1.8% | 1.8% |
| | | faeces | digestive tract contents | 16 | 1.6% | 1.6% |
| | | lake sediment | permafrost | 9 | 0.9% | 0.9% |
| | | Unknown | bone | 9 | 0.9% | 0.9% |
| | | birch pitch | saliva | 7 | 0.7% | 0.7% |
| | | intestine | skin | 7 | 0.7% | 0.7% |

**Category:** Seen labels were present in training; Unseen labels were not in training. **Seen - Correct:** Samples correctly classified with labels seen during training. **Seen - Wrong:** Top 10 most frequent misclassification patterns for seen labels (True Label → Predicted Label). **Unseen:** Top 10 most frequent predictions for unseen labels (model maps to semantically similar seen classes). **% Task:** Percentage relative to total samples for that task in validation set. **% Total:** Percentage relative to total validation samples across all tasks. Total validation samples: 987 (unique sample IDs across 4 tasks = 3948 predictions).

Supplementary Table 8: Parameters and computational resources used to build the unitig feature matrix with muset

| Category | Parameter | Value |
|---|---|---|
| Input | Samples | 3,070 |
| | Data source | Logan unitigs |
| k-mer Filtering | k-mer size | 31 |
| | Minimum abundance | 2 |
| | Minimizer size | 15 |
| | Min. fraction present (-F) | 0.1 |
| | Max. fraction absent (-f) | 0.1 |
| | Initial k-mers | 436,456,132,058 |
| | Retained k-mers | 18,953,119 |
| Unitig Assembly | Minimum length | 61 bp |
| | Total unitigs assembled | 2,755,241 |
| | Final unitigs | 107,480 |
| Computational | Tool | muset v0.6.1 |
| | CPU cores | 32 |
| | Memory | 500 GB |
| | Runtime | 3d 4h 45m |

**Columns**: *Category*—group of related parameters (Input: raw data; k-mer Filtering: counting and filtering settings; Unitig Assembly: graph-assembly settings; Computational: hardware and runtime). *Parameter*—specific setting or measured quantity. *Value*—value used or observed during matrix construction. k-mer size 31 bp with minimum abundance 2 removes sequencing errors. Fraction filters (-F/-f) retain unitigs present in at least 10% of samples and absent from at most 90% of samples, thereby keeping only informative features. Final unitigs (107,480) are those passing all filters and assembled into sequences $\geq$61 bp.

Supplementary Table 9: Top 15 BioProjects contributing the most prediction errors (train + test + validation combined, all four tasks).

| BioProject | Study | Train err. | Test err. | Val err. | Total | Err. (%)[a] | Cum. (%)[b] | Error type[c] | Main errors[d] |
|---|---|---|---|---|---|---|---|---|---|
| PRJNA994900 | Kirdok2024 | – | – | 210 | 210 | 75.0 | 13.4 | Absent class (Mat.) | birch pitch→sediment (Mat.); ancient→modern (ST) |
| PRJEB41240 | SeguinOrlando2021 | 149 | 21 | 12 | 182 | 19.1 | 25.0 | Genuine confusion | bone→tooth/sediment (Mat.); *H. sapiens*→n.a. (SH) |
| PRJNA354503 | Philips2017 | 112 | 21 | 16 | 149 | 22.6 | 34.5 | Genuine confusion | oral→skel. tissue (CT); tooth→bone (Mat.) |
| PRJEB64128 | Jackson2024 | – | – | 122 | 122 | 41.8 | 42.3 | Genuine confusion | tooth→dent. calc. (Mat.); oral→skel. tissue (CT) |
| PRJEB34569 | FellowsYates2021 | 59 | 6 | 42 | 107 | 13.3 | 49.1 | Taxonomic granularity (SH) | *G. b. beringei*→*Gorilla* sp.; *P. t. schweinfurthii*→*P. troglodytes* |
| PRJEB61887 | Velsko2024 | – | – | 99 | 99 | 20.3 | 55.4 | Genuine confusion | oral→skel. tissue (CT); bone→tooth (Mat.) |
| PRJEB33848 | Neukamm2020 | 27 | 7 | 32 | 66 | 10.7 | 59.6 | Genuine confusion; Absent class (Mat.) | soft tis.→skel. tis. (CT); Unknown→bone (Mat.) |
| PRJEB49638 | Moraitou2022 | 18 | 10 | 32 | 60 | 15.8 | 63.5 | Taxonomic granularity (SH) | *G. b. graueri*→*Gorilla* sp.; *G. gorilla gorilla*→*Gorilla* sp. |
| PRJNA1211513 | Schreiber2025 | – | – | 48 | 48 | 25.0 | 66.5 | Label granularity (Mat.) | marine sed.→sediment (Mat.) |
| PRJEB80877 | vonHippel2025 | – | – | 46 | 46 | 25.0 | 69.5 | Label granularity (Mat.) | lake sed.→sediment (Mat.) |
| PRJNA836960 | Madison2023 | – | – | 35 | 35 | 79.5 | 71.7 | Absent class (Mat., SH) | intestine→skin (Mat.); gut→soft tissue (CT) |
| PRJEB67998 | Bozzi2024 | – | – | 34 | 34 | 44.7 | 73.9 | Genuine confusion | tooth→bone/skin (Mat.); skel. tis.→soft tis. (CT) |
| PRJNA1056444 | Austin2024 | – | – | 28 | 28 | 17.9 | 75.6 | Genuine confusion | *H. sapiens*→*Pan troglodytes* (SH) |
| PRJEB11419 | – | – | – | 26 | 26 | 65.0 | 77.3 | Label granularity (Mat.) | faeces→dig. tract (Mat.); modern→ancient (ST) |
| PRJNA48479 | – | – | – | 25 | 25 | 32.9 | 78.9 | Genuine confusion | modern→ancient (ST); oral→skel. tissue (CT) |

**Columns**: *BioProject*—NCBI BioProject accession. *Study*—first-author and year of the associated publication. *Train/Test/Val err.*—number of wrong predictions (across all four tasks) in each dataset split; '–' means the BioProject has no samples in that split. *Total*—sum of wrong predictions across all three splits. [a] *Err. (%)*: wrong predictions as a percentage of all predictions made for that BioProject (each sample contributes four predictions, one per task). [b] *Cum. (%)*: running total of errors as a percentage of all errors across every qualifying BioProject (≥5 samples in test + validation); shows how concentrated errors are in a few studies. [c] *Error type*: four categories are used. *Absent class*: the true label never appeared in training. *Label granularity*: the true label is absent from training but is a biological subtype of a training class (e.g. `lake sediment` ⊂ `sediment`); the model predicts the correct coarser label. *Taxonomic granularity*: the true host is a subspecies whose parent taxon is the training label (e.g. *Gorilla beringei beringei* → *Gorilla* sp. in training). *Genuine confusion*: all true labels were seen in training; the model still predicts the wrong class. [d] *Main errors*: top true → predicted label pairs by frequency. Task abbreviations: ST = Sample Type, CT = Community Type, SH = Sample Host, Mat. = Material.

### 2 Supplementary Figures

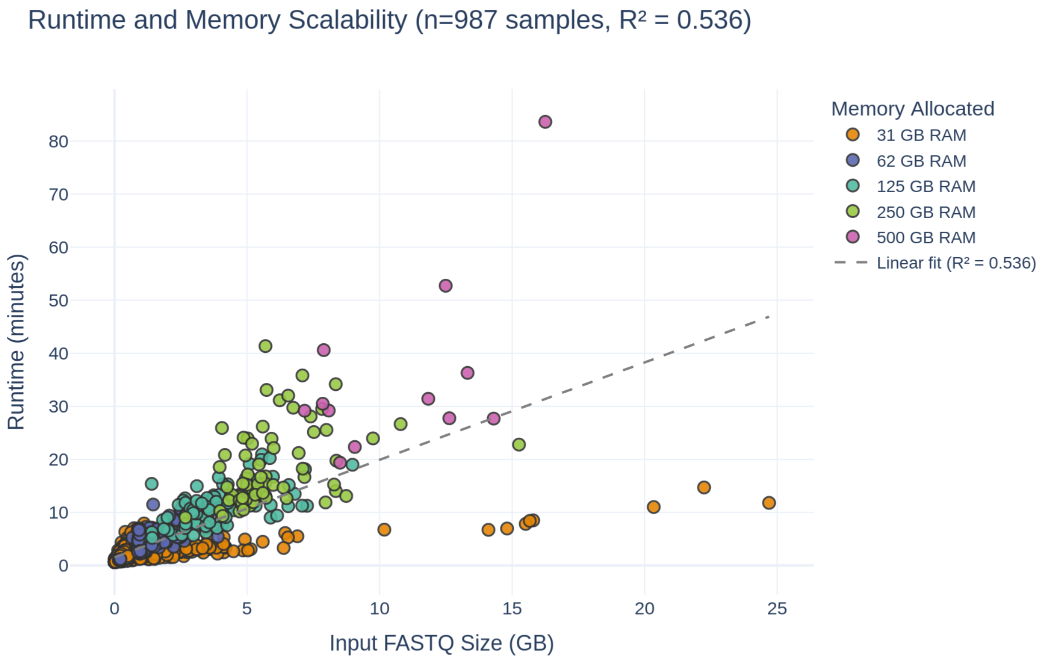

Supplementary Figure 1: **Runtime and memory scalability analysis.** Scatter plot showing the relationship between input FASTQ file size (GB) and runtime (minutes) for DIANA predictions on the validation set. Points are colored by RAM allocation (32-512 GB). Linear regression fit (grey dashed line) demonstrates strong linear scaling ($R^2 > 0.536$), indicating predictable computational requirements proportional to input data size. All measurements were obtained from the validation set. Each point represents one successfully processed sample.

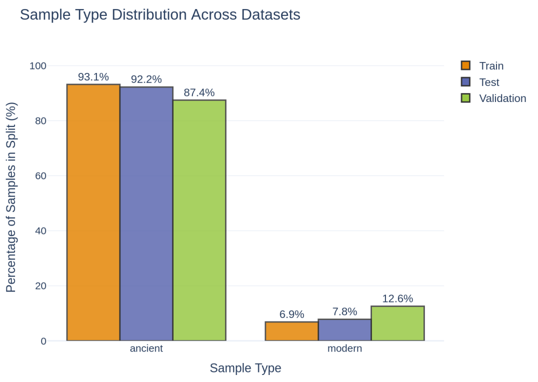

(a) Sample Type Distribution

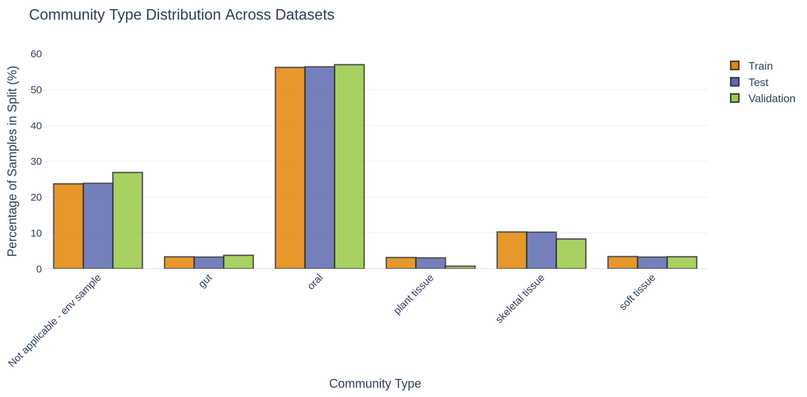

(b) Community Type Distribution

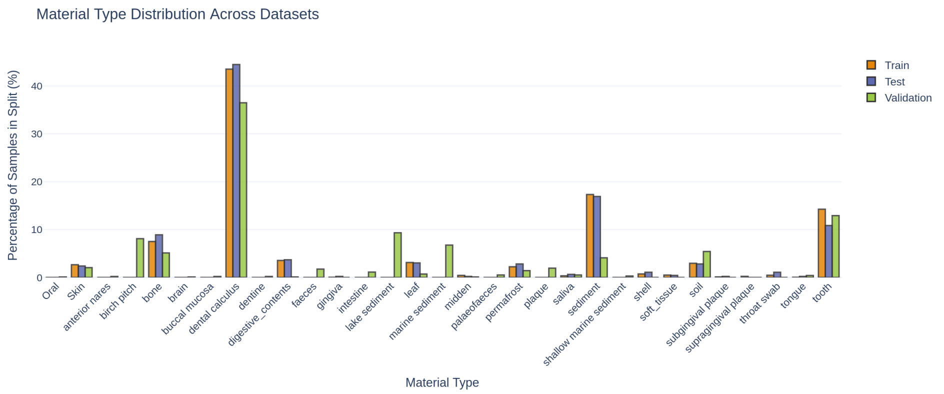

(c) Material Distribution

Supplementary Figure 2: **Distribution comparison across train, test, and validation splits.** (a) Sample type shows ancient vs. modern metagenome distribution, with balanced representation across all train and test. (b) Community type shows 6 classes. (c) Material shows 13 substrate types in training, with validation including 7 additional unseen types. *(Figure continued on next page)*

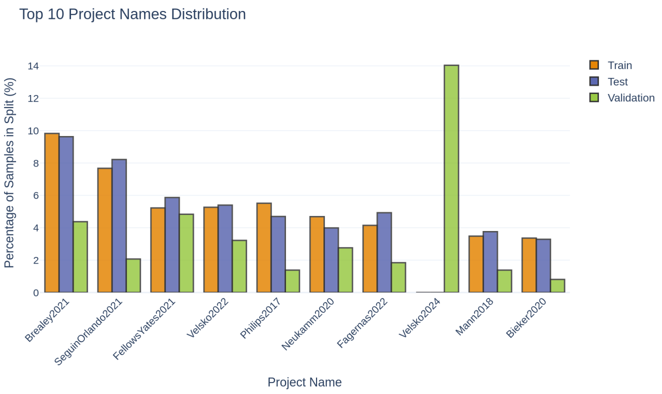

(d) Project Distribution

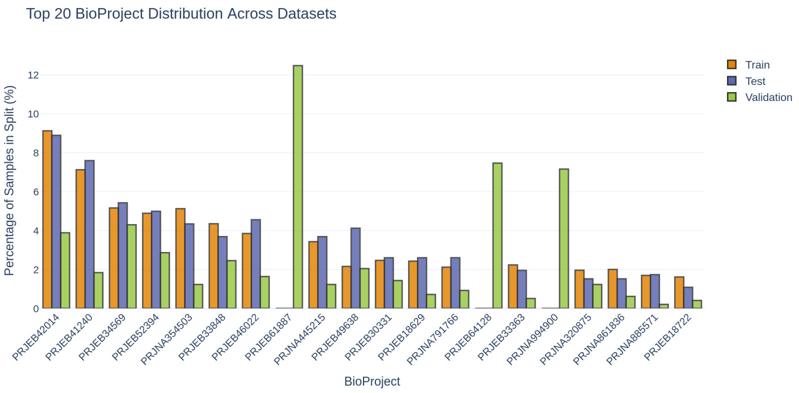

(e) BioProject Distribution

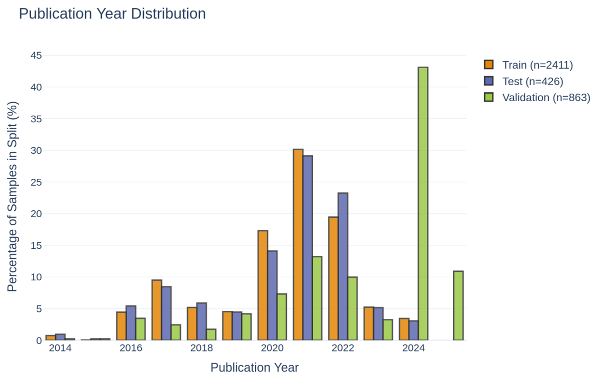

(f) Publication Year Distribution

Supplementary Figure 2: **Distribution comparison across train, test, and validation splits (continued).** (d) Project distribution across major sequencing initiatives. (e) BioProject distribution across datasets. (f) Publication year distribution demonstrates temporal coverage from 2014-2025. *(Figure continued on next page)*

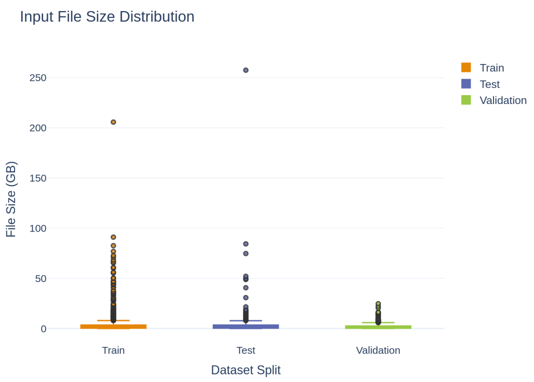

(g) File Size Distribution

Supplementary Figure 2: **Distribution comparison across train, test, and validation splits (continued).** (g) File size distribution (GB) shows varying input file sizes across datasets. Bar plots showing the distribution of samples across different metadata categories for the training set (n=2,597), held-out test set (n=461), and external validation set (n=987).

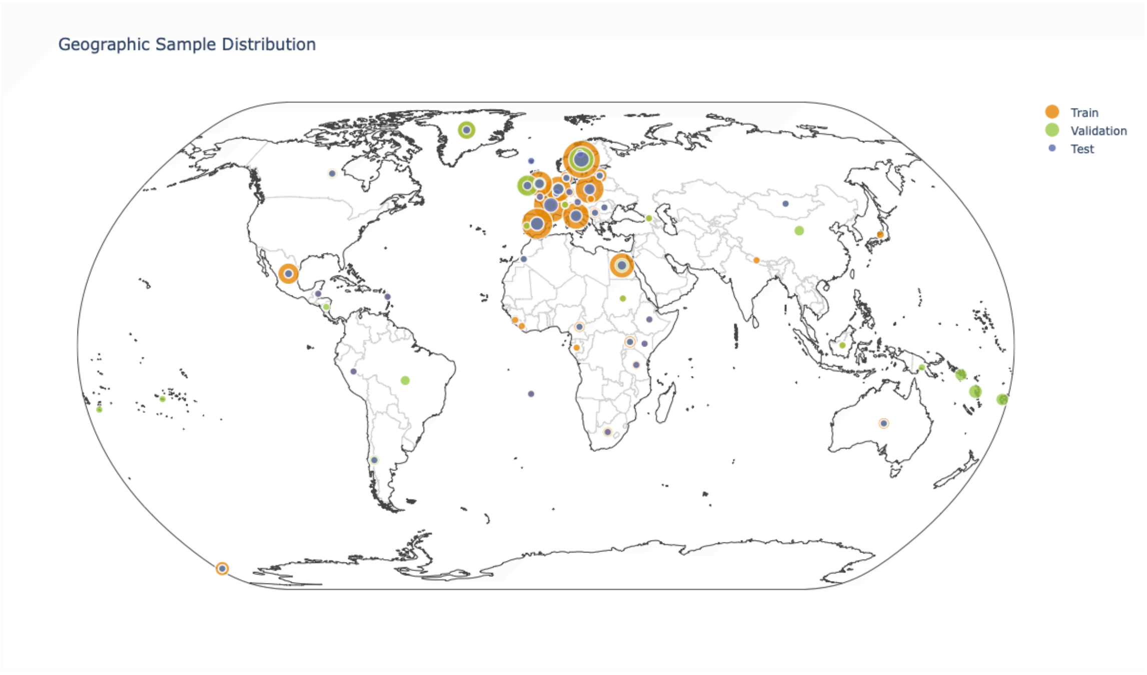

Supplementary Figure 3: **Geographic distribution of samples by dataset.** World map showing global sampling coverage with points colored by dataset and sized by sample count. Training (orange, n=2,597) and test (blue, n=461) samples were predominantly from Europe and North America. Validation samples (green, n=987). The sequencing bias that exists in the field is going to be reflected in the performance of .

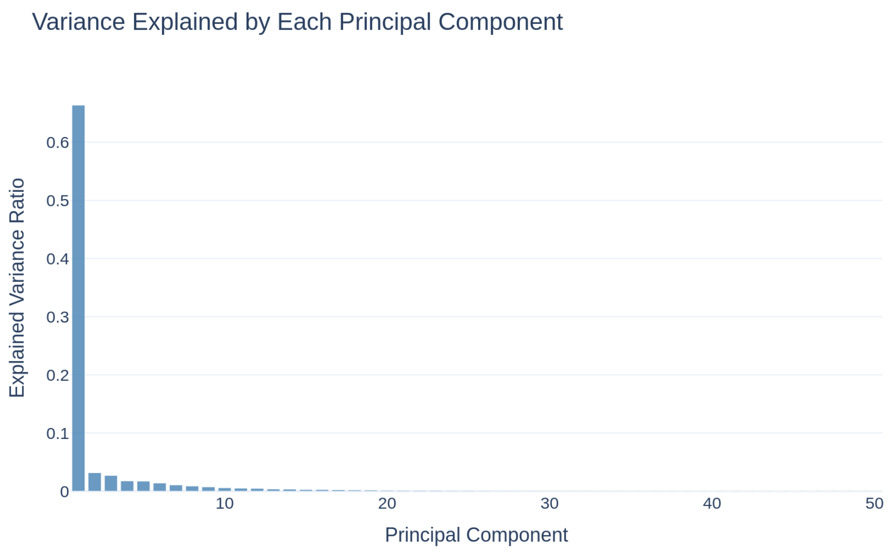

(a) Variance explained per component

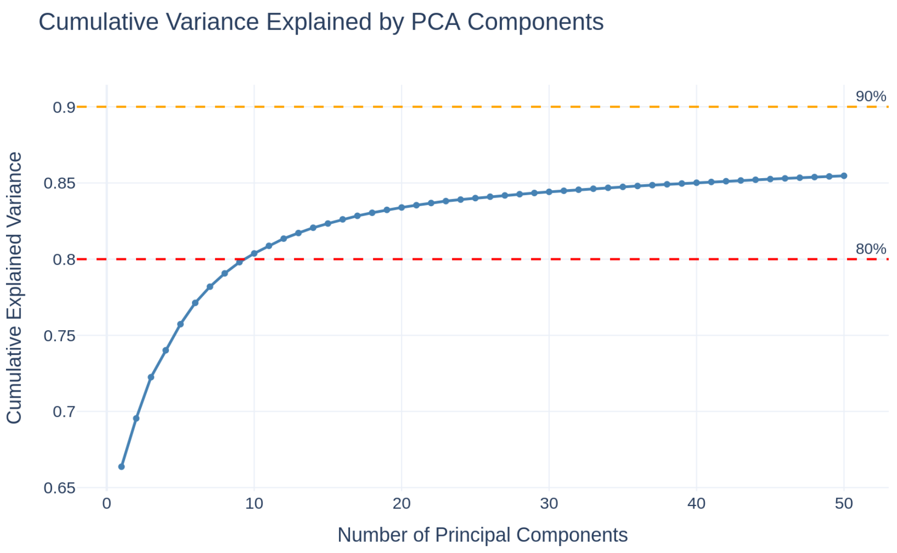

(b) Cumulative variance explained

Supplementary Figure 4: **PCA variance explained by the unitig feature matrix.** PCA computed on training and test samples (n=3,058: 2,597 train + 461 test) using 107,480 unitig features. **(a)** Variance explained by each of the first 50 principal components. PC1 alone accounts for 66.4% of total variance, showing a dominant first component with a sharp drop to PC2 (3.2%) and a gradual, long tail thereafter. **(b)** Cumulative variance explained. The first two components together capture approximately 70% of total variance; the 80% threshold is reached at ~10 components, and 90% is not reached even at 50 components (~86%). Although PC1 captures substantial variance, the remaining variance is spread across many components.

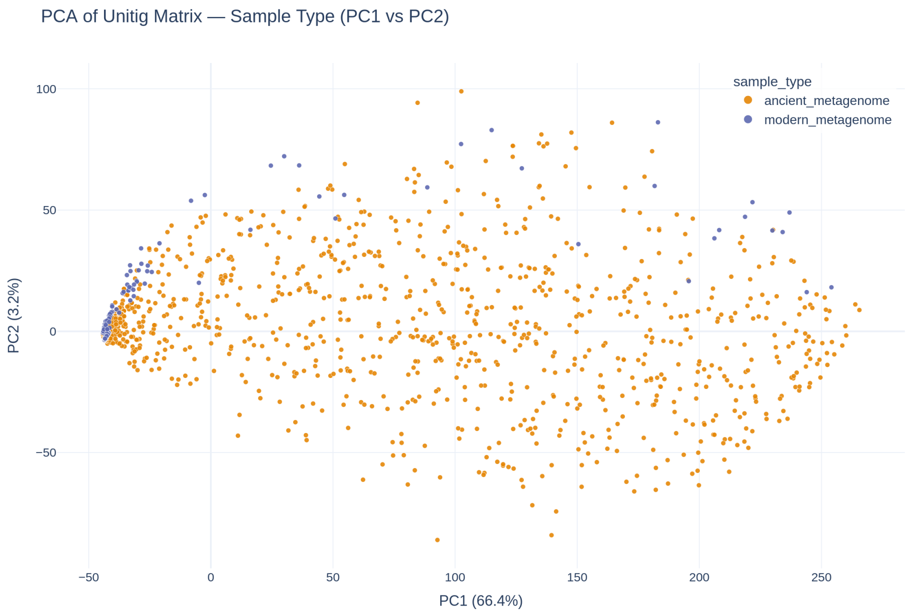

(a) Sample Type

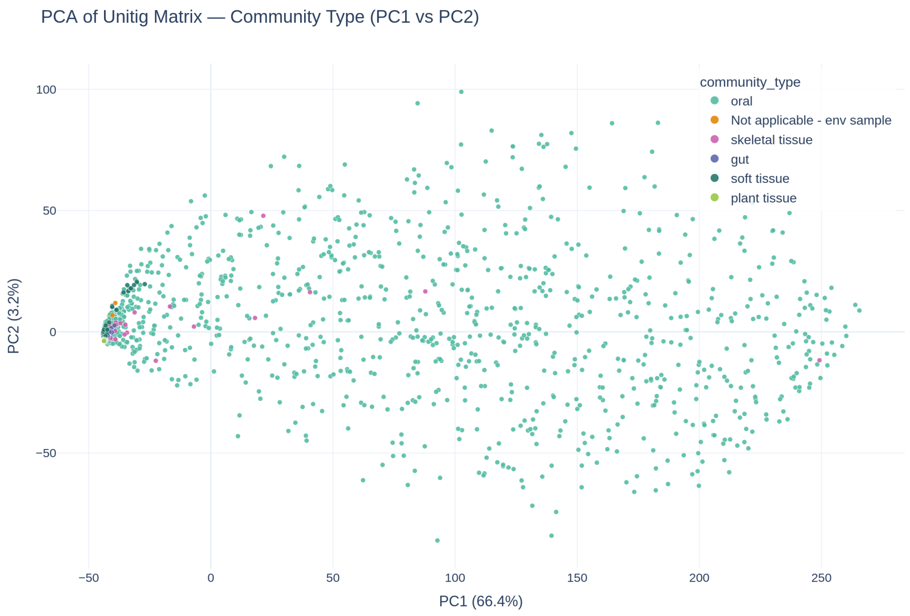

(b) Community Type

Supplementary Figure 5: **PCA projections colored by classification task labels.** 2D projections of training and test samples (n=3,058 total: 2,597 train + 461 test) onto PC1 and PC2, colored by classification task labels. There is no clear separation of the distinct classes for any of the 4 classification tasks attempted. (a) Sample type (2 classes: ancient vs. modern). (b) Community type (6 categories). *(Figure continued on next page)*

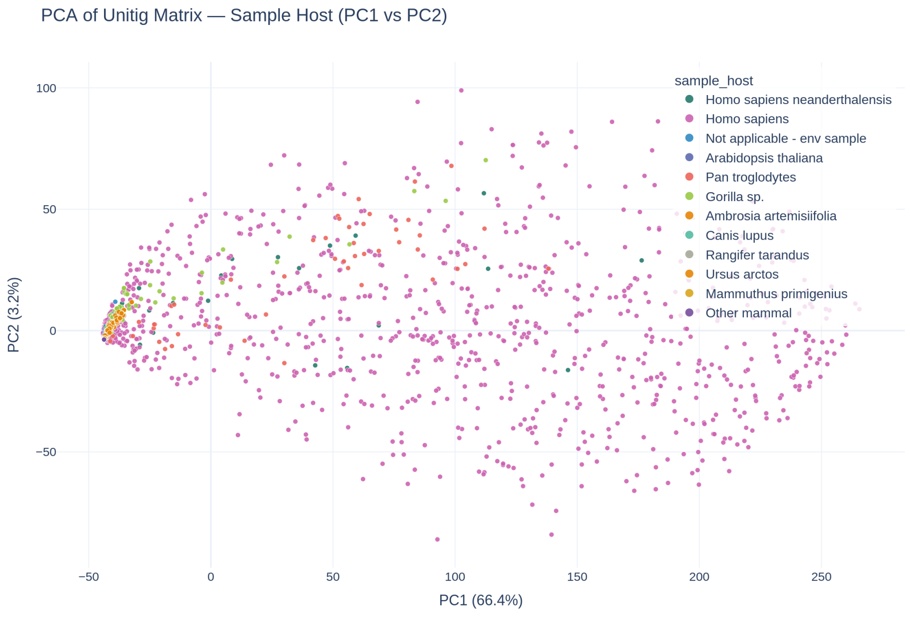

(c) Sample Host

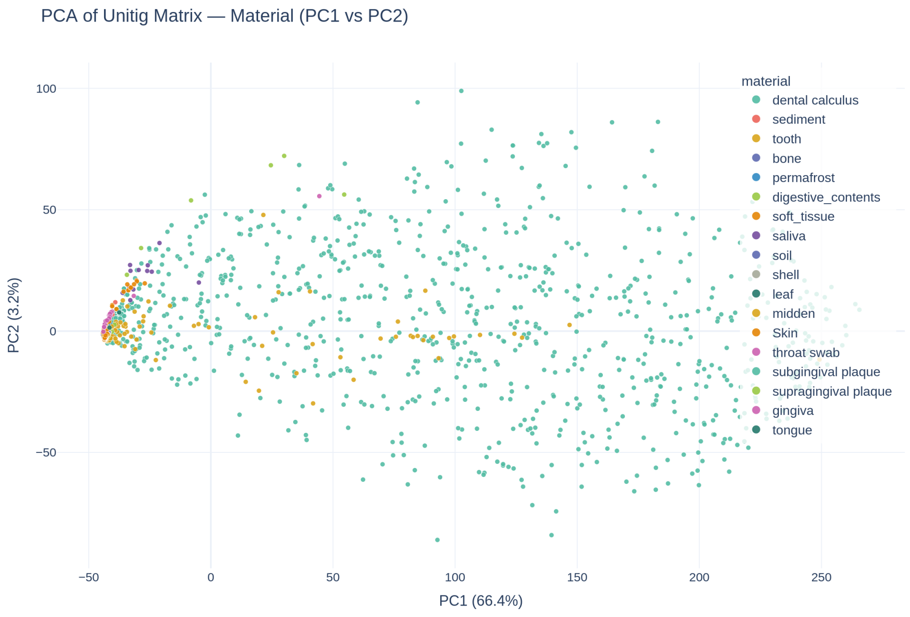

(d) Material

Supplementary Figure 5: **PCA projections colored by classification task labels (continued).** (c) Sample host (12 species in training). (d) Material (13 types in training).

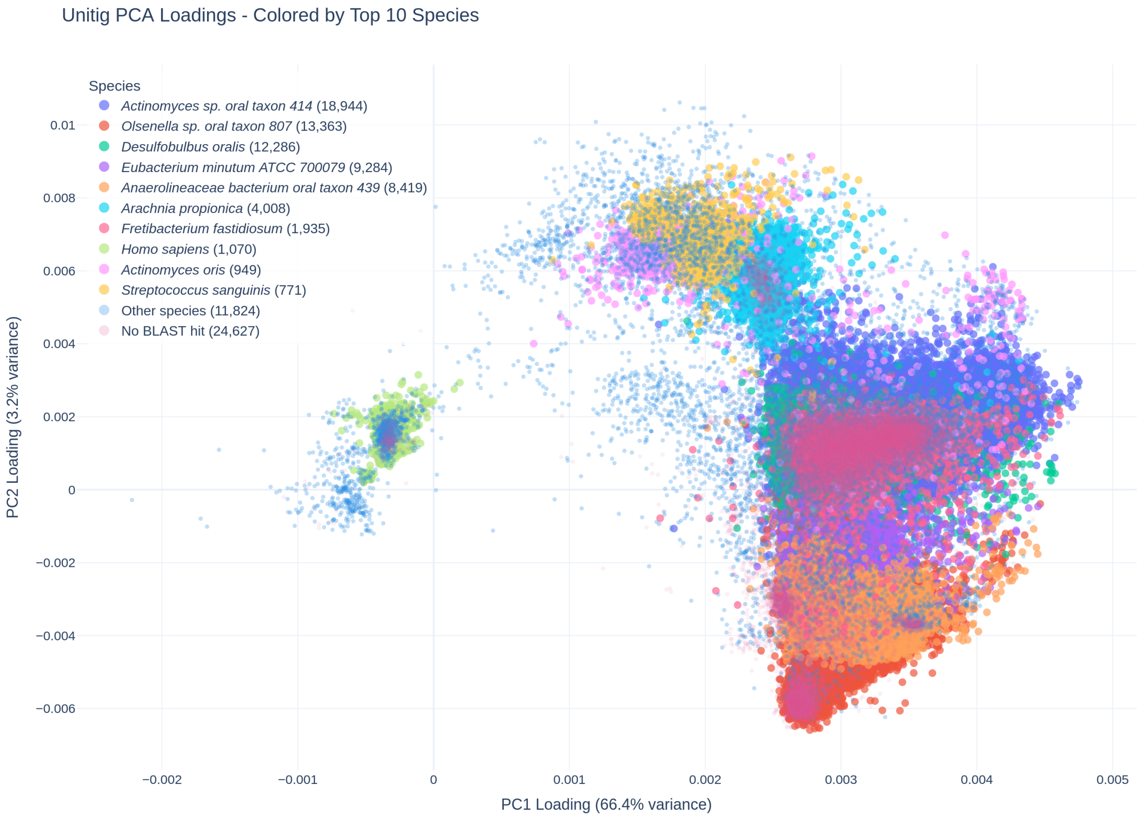

Supplementary Figure 6: **Unitig PCA loadings colored by taxonomic species.** Scatter plot showing PC1 and PC2 loadings for all 107,480 unitig features, colored by the top 20 most abundant species from BLAST annotations. Points represent individual unitigs, with position indicating their contribution to each principal component. Species with the largest numbers of annotated unitigs are shown in distinct colours, while other species and unannotated unitigs are displayed in light grey for context. Distinct clustering of unitigs by species demonstrates that the principal components capture taxonomically meaningful variation, including a clear separation between *Homo sapiens* and most of the bacterial sequences. Interestingly, a small group of non-human unitigs overlaps with the *Homo sapiens* cluster, possibly reflecting compositional similarity, misclassification, or shared genomic regions between host and microbiome sequences.

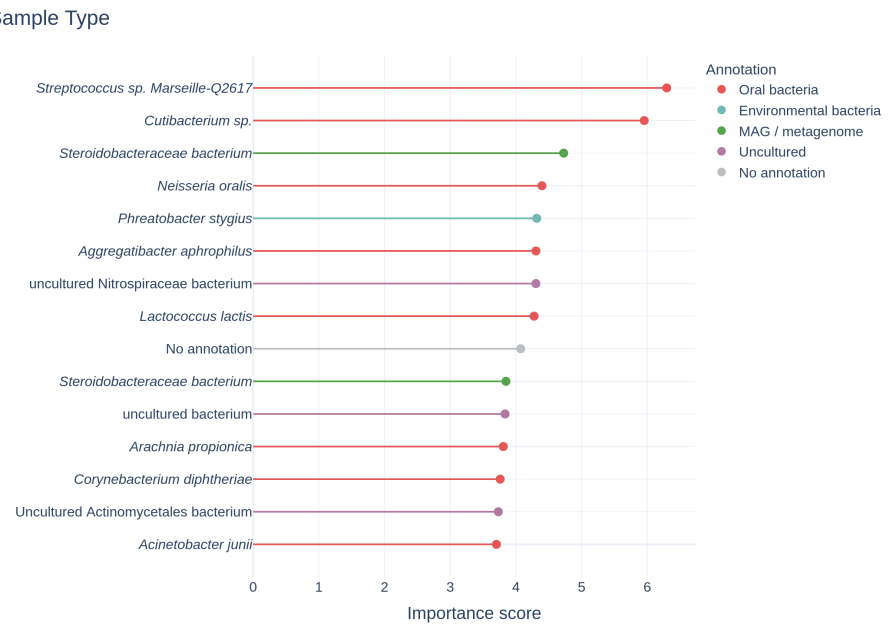

(a) Sample Type

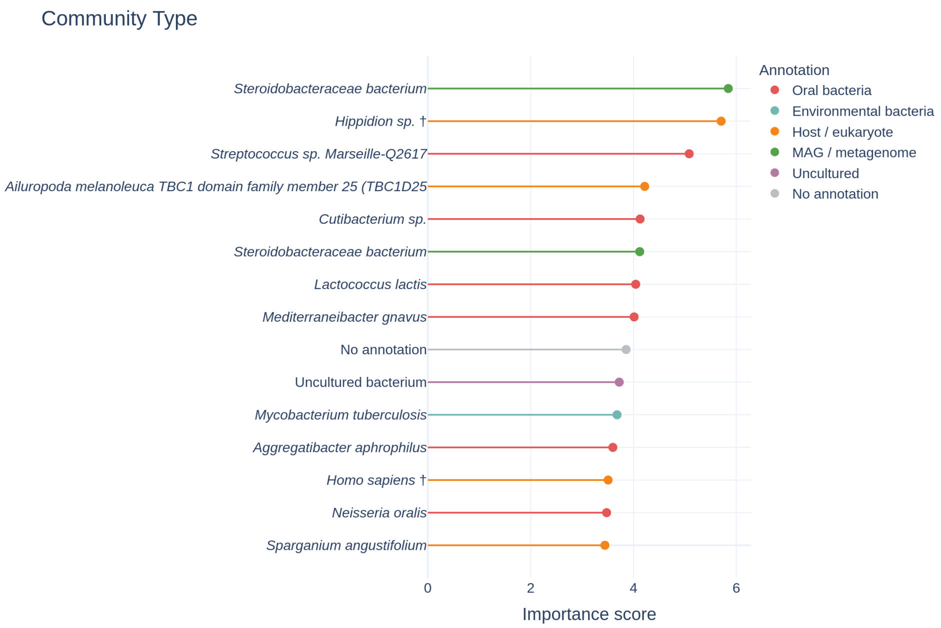

(b) Community Type

Supplementary Figure 7: **Top-15 most important unitig features per classification task, annotated by biological origin.** Lollipop plots showing the 15 highest-importance unitig features for each task. (a) Sample type. (b) Community type. Feature importance is the mean gradient-based score across training epochs. Each feature is labelled with its species-level BLAST hit against the NCBI nt database (mean query coverage 98.3%); the colour encodes the biological category of that annotation. *(Figure continued on next page)*

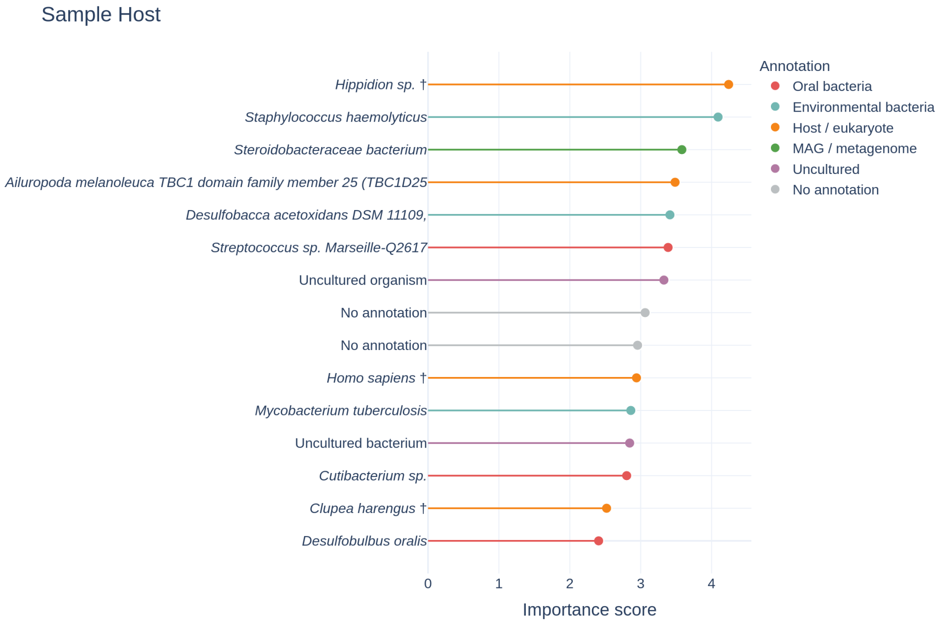

(c) Sample Host

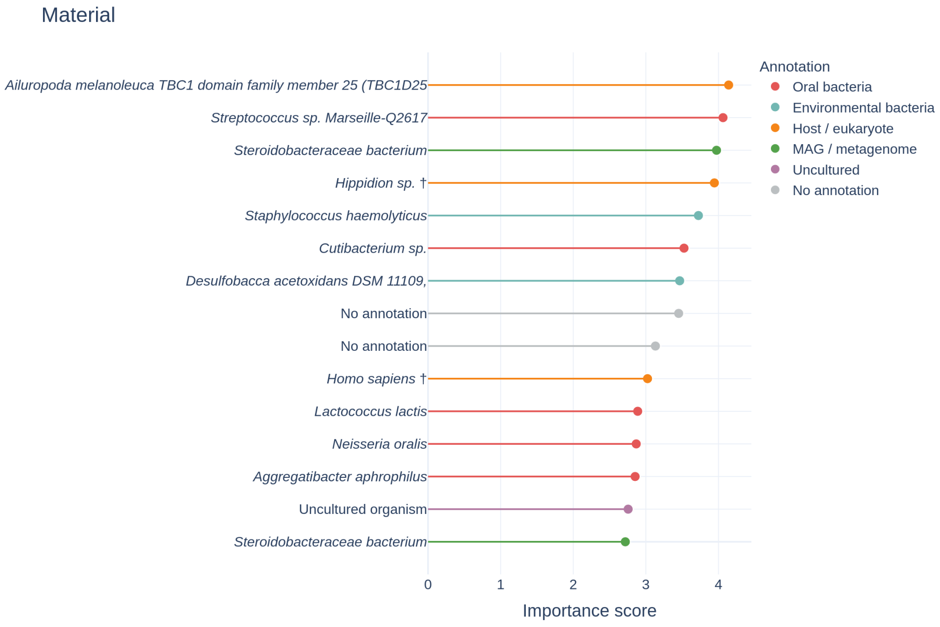

(d) Material

Supplementary Figure 7: **Top-15 most important unitig features per classification task (continued).** (c) Sample host. (d) Material. Features marked † could not be matched by BLAST and were instead annotated via the Logan SRA genomic index (best hit by percent identity). Features labelled *No annotation* are simple tandem repeats (e.g. $(AGT)_n$, $(CACT)_n$) that were masked in both databases.

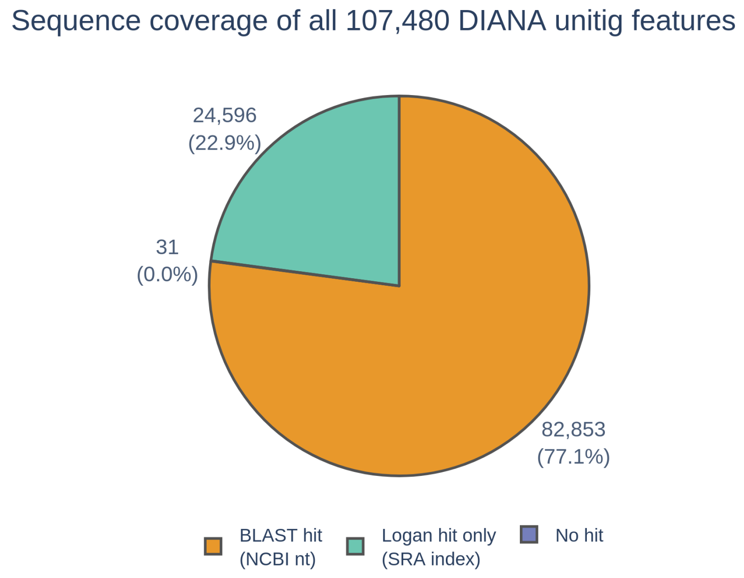

Supplementary Figure 8: **Taxonomic annotation coverage of DIANA unitig features.** Proportion of all 107,480 unitig features annotated by a two-stage strategy: BLAST against NCBI nt (77.1%, n=82,853; mean query coverage 98.3%), followed by Logan search against the SRA genomic index for unannotated unitigs (22.9%, n=24,596; mean query coverage 99.7%). Only 31 features (0.03%) remained unannotated after both stages.

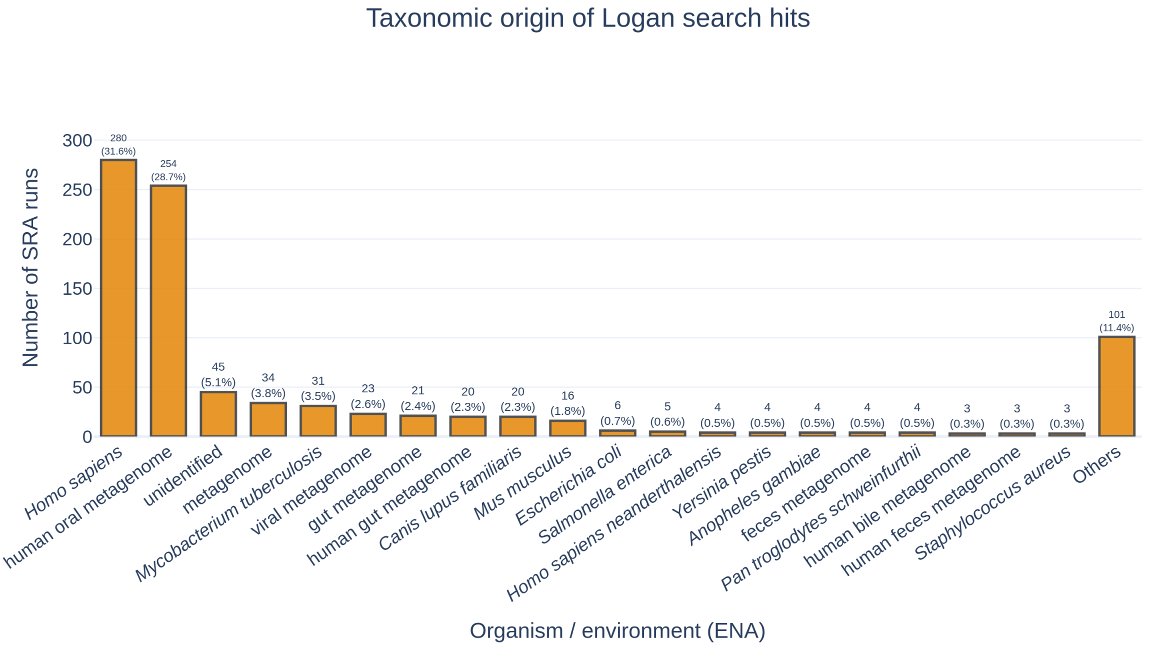

Supplementary Figure 9: **Unitigs found in Logan-search.** Bar chart showing the top 20 scientific names assigned to the 885 SRA runs that provided the best Logan hit for unitigs absent from NCBI nt (n=24,596 unitigs).

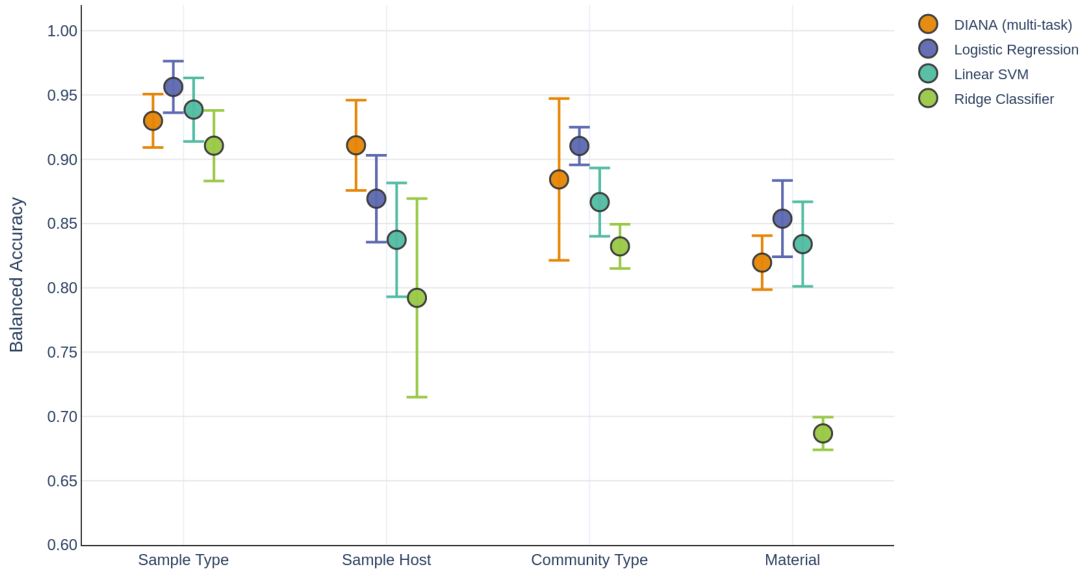

Supplementary Figure 10: **DIANA outperforms single-task linear baselines on the training set.** Five-fold cross-validation balanced accuracy (mean ± std) on the training set (n=2,597) for DIANA and three class-balanced linear classifiers trained independently per task: Logistic Regression, Linear SVM, and Ridge Classifier. Tasks are ordered by decreasing DIANA performance (left to right). DIANA achieves the highest balanced accuracy on three of four tasks (Sample Type: 0.969, Community Type: 0.918, Sample Host: 0.904) using a single jointly-trained model, while baselines require four independent models. On the 13-class Material task, linear baselines marginally outperform DIANA on average, though overlap within DIANA's cross-validation variance (±0.095) precludes a definitive conclusion.

### 3 Supplementary Methods

#### 3.1 Adapter contamination check

We compiled 39 unique Illumina adapter sequences from two standard adapter lists available at `https://github.com/CamilaDuitama/DIANA/tree/paper/data/adapters/` and mapped them against all 107,480 unitig sequences (`unitigs.fa`; Zenodo `https://zenodo.org/records/18496961`) using Bowtie 2 v2.4.5:

```
bowtie2-build unitigs.fa unitigs_bt2_index
bowtie2 -x unitigs_bt2_index -f -U adapters.fasta \
        --very-sensitive --no-unal -S adapter_hits.sam
```

38 of 39 adapters produced zero alignments. The single partial match (13 bp TruSeq core motif `AGATCGGAAGAGC`, $MAPQ = 0$) was not among the top discriminative features for any task, and the exact sequence is absent from all unitigs.
